# Biosynthesis system of Synechan, a sulfated exopolysaccharide, in the model cyanobacterium *Synechocystis* sp. PCC 6803

**DOI:** 10.1101/2021.02.01.429190

**Authors:** Kaisei Maeda, Yukiko Okuda, Gen Enomoto, Satoru Watanabe, Masahiko Ikeuchi

## Abstract

Extracellular polysaccharides of bacteria contribute to biofilm formation, stress tolerance, and infectivity. Cyanobacteria, the oxygenic photoautotrophic bacteria, uniquely and widely have sulfated extracellular polysaccharides and they may utilize the polysaccharides for survival in nature. In addition, sulfated polysaccharides of cyanobacteria and other organisms have been focused as beneficial biomaterial. However, very little is known about their biosynthesis machinery and function in cyanobacteria. Here we found that the model cyanobacterium, *Synechocystis* sp. PCC 6803, formed bloom-like cell aggregates using sulfated extracellular polysaccharides (designated as synechan) and identified whole set of genes responsible for synechan biosynthesis and its transcriptional regulation, thereby suggesting a model for the synechan biosynthesis apparatus. Because similar genes are found in many cyanobacterial genomes with wide variation, our findings may lead elucidation of various sulfated polysaccharides, their functions, and their potential application in biotechnology.

## Introduction

Bacterial extracellular polysaccharides establish biofilms for nutrient supply and stress avoidance, and they sometimes support cellular activities such as motility and infectivity (*Woodward and Naismith, 2016*). Generally, the polysaccharide chains consist of a few types of sugars (with or without chemical modifications) and are anchored on cells (capsular polysaccharides, CPS) or exist as nonanchored exopolysaccharides (EPS). Nonetheless, their molecular structures vary greatly, e.g., branching schemes, sugar constituents, and modifications, and thus their physical properties also vary. Bacterial extracellular polysaccharides and lipopolysaccharides are produced and exported via three distinct pathways: the Wzx/Wzy-dependent pathway, ABC-dependent pathway, and synthase-dependent pathway (*Schmid et al., 2015*). Every bacterium can produce several extracellular polysaccharides, and production often depends on environmental conditions. Some extracellular polysaccharides have been appropriated for use as biopolymers for food, cosmetics, medicine (*Freitas et al., 2014, Lapasin and Pricl, 1995*).

Cyanobacteria, the oxygenic photoautotrophic bacteria that inhabit almost every ecosystem on Earth, contribute to the global photosynthetic production (*Flombaum et al., 2013, Mangan et al., 2016*). Cyanobacteria produce various extracellular polysaccharides to form colonies, which are planktonic or attached on solid surfaces, likely to stay in a phototrophic niche in nature (*De Philippis and Vincenzini, 1998*). A notable example is the water bloom, a dense population of cyanobacterial cells that floats on the water surface and often produces cyanotoxins and extracellular polysaccharides (*Huisman et al., 2018*). The extracellular polysaccharides are also important for photosynthetic production of cyanobacteria and their application (*Kumar et al., 2018*). However, very little is known about their biosynthesis except for extracellular cellulose. A thermophilic cyanobacterium (*Thermosynechococcus vulcanus*) accumulates cellulose to form cell aggregation (*Kawano et al., 2011*). This cellulose is produced by cellulose synthase with unique tripartite system and regulations (*Enomoto et al., 2015, Maeda et al., 2018*). In the cyanobacterial genomes, there are still many putative genes for extracellular polysaccharide biosynthesis.

Uniquely, many cyanobacterial extracellular polysaccharides are sulfated, i.e., as a sugar modification (*Pereira et al., 2009*). Sulfated polysaccharides are also produced by animals (as glycosaminoglycan in the extracellular matrix such as heparan sulfate) and algae (as cell-wall components such as carrageenan) but are scarcely known in other bacteria or plants (*Ghosh et al., 2009*). Major examples of cyanobacterial sulfated polysaccharides are spirulan from *Arthrospira platensis* (vernacular name, “Spirulina”), sacran from *Aphanothece sacrum* (vernacular name, “Suizenji-Nori”) and cyanoflan from *Cyanothece* sp. CCY 0110 (*Mota et al., 2020, Mouhim et al., 1993, Okajima et al., 2008*). These sulfated polysaccharides are used for formation of colony and biofilm and may be functionally relevant to the ecology of cyanobacteria (*Fujishiro et al., 2004*). In addition, the bioactivities (antiviral, antitumor, and anti-inflammatory) of sulfated polysaccharides from cyanobacteria were reported, too (*Flores et al., 2019a, Hayashi et al., 1996, Ngatu et al., 2012*). However, very little is known about their biosynthesis machinery and physiological functions. On the other hand, biosynthesis and modification of animal sulfated polysaccharides have been extensively studied because of their importance to tissue protection, tissue development, and immunity (*Karamanos et al., 2018, Sasisekharan et al., 2006*) and potential applications in healthy foods, biomaterials, and medicines (*Jiao et al., 2011, Wardrop and Keeling, 2008*).

The poor understanding about cyanobacterial sulfated polysaccharide biosynthesis is probably due to low or no accumulation of sulfated polysaccharides in typical model species (*Pereira et al., 2009*). More than three decades ago, Panoff et al. reported sulfated polysaccharides in two related model cyanobacteria, *Synechocystis* sp. PCC 6803 and *Synechocystis* sp. PCC 6714 (hereafter *Synechocystis* 6803 and *Synechocystis* 6714) (*Panoff et al., 1988*). Recently, Flores et al. confirmed sulfated polysaccharides and reported their enhanced accumulation in a sigma factor *sigF* mutant for global cell surface regulation in *Synechocystis* 6803 (*Flores et al., 2019b*). In parallel, several papers have studied genes that could be involved in extracellular polysaccharides biosynthesis in *Synechocystis* 6803, but no clear results were obtained about the sulfated polysaccharide (*Fisher et al., 2013, Foster et al., 2009, Jittawuttipoka et al., 2013, Pereira et al., 2019*). Here we found that a motile substrain of *Synechocystis* 6803 showed bloom-like cell aggregation and sulfated EPS production, but a non-motile substrain (a standard substrain for photosynthesis study) did not. By gene disruption and overexpression, we first identified a whole set of genes responsible for sulfated EPS biosynthesis and its regulatory system, opening the way to engineering of their production.

## Results

### *Bloom formation and EPS accumulation in* Synechocystis *sp. PCC 6803*

We fortuitously found that a motile substrain of *Synechocystis* 6803 produces EPS and forms floating cell aggregates resembling a typical cyanobacterial bloom. We established a two-step culture regime (2-day bubbling culture and subsequent standing culture without bubbling under continuous light) for reproducible formation of bloom-like aggregates (Fig. 1A, B). The first (bubbling) step allows for cell propagation and EPS production, whereas the second (standing) step allows for heavy-cell aggregation and flotation, even though *Synechocystis* 6803 does not possess genes for intracellular gas vesicles (*Harke et al., 2016*). In *Synechocystis* 6803, cell flotation accompanying the generation of extracellular gas bubbles was suppressed by inactivation of photosynthesis (Fig. 1C), suggesting that gas derived from photosynthesis drives the upward movement of cells embedded in viscous EPS. The non-motile, glucose-tolerant substrain—commonly used for photosynthesis research—did not aggregate or float.

**Figure 1.**
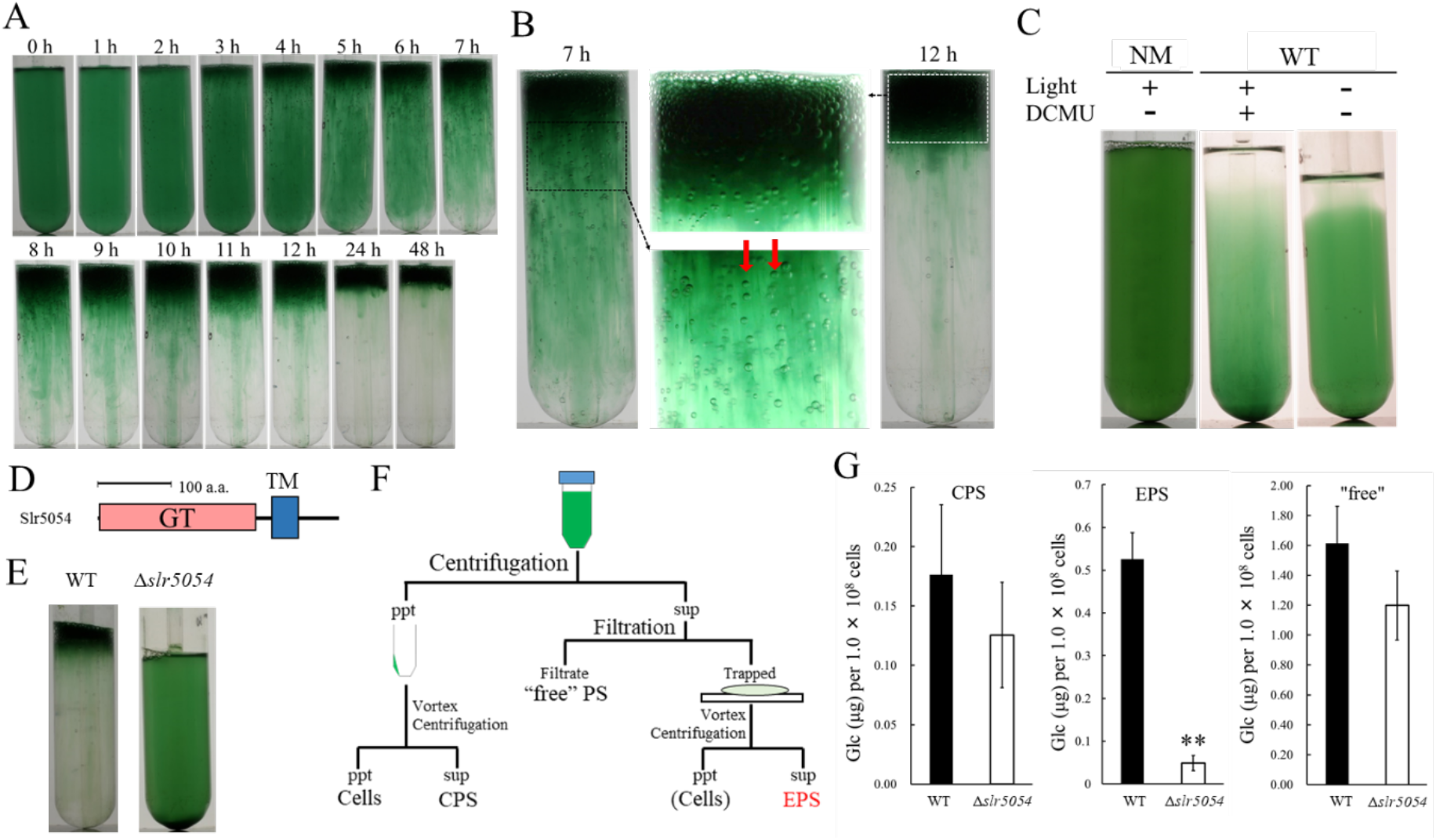
Bloom formation and EPS isolation. **A**, Time course of bloom formation by WT *Synechocystis* 6803 during the second step of culture. Extracellular gas bubbles are formed and trapped in viscous EPS (~1 h). Green vertical columns with bubbles become apparent at 4 h. Those trapped gas bubbles slowly rise together with the viscous columns. **B**, Enlarged images showing gas bubbles trapped in EPS. Vertically aligned bubbles are indicated by red arrows. **C**, Lack of bloom formation in the non-motile substrain (NM) or WT with or without light and the photosynthesis inhibitor DCMU at 48 h of the second step of culture. **D**, Domain architecture of Slr5054. GT, glycosyltransferase domain; TM, transmembrane region. **E**, Lack of bloom formation in Δ*slr5054* after standing culture for 48 h. **F**, Isolation of EPS from the first step of culture. Cells and CPS were removed from the culture by centrifugation, and EPS in the supernatant was separated from “free” polysaccharide (PS) by membrane filtration followed by a second centrifugation to remove residual cells. CPS was collected from the cell pellet after vortexing and centrifugation. **G**, Sugar content of fractions from WT and Δ*slr5054*. Error bars represent SD (n = 3, \*\**P* < 0.005).

We first isolated crude EPS from the bloomed culture by membrane filtration (Fig. S1A). The crude EPS consisted of polysaccharide but very little protein or nucleic acid, and its abundance remained unchanged during the second culture step (Fig. S1B). As a common feature of diverse EPS biosynthesis systems in bacteria, membrane-bound glycosyltransferases are particularly important (*Schmid et al., 2015*). So, we screened such glycosyltransferase genes by disruption and revealed that *slr5054* is essential for bloom formation (Fig. 1D, E and Fig. S2). The EPS preparation was improved by removing cells before filtration to avoid cell-associated polysaccharides such as CPS (Fig. 1F). Δ*slr5054* lacked most of the EPS present in the wild type (WT), whereas the CPS and free polysaccharide fractions were similar in the WT and Δ*slr5054* (Fig. 1G). Then we performed Alcian blue staining to examine the acidity of the EPS (Fig. S3). Generally, sulfated polysaccharides are stained at pH 0.5 condition, while acidic polysaccharides, which contain sulfate groups and/or carboxylate groups (such as uronic acids and carboxylate modification) are stained at pH 2.5 condition (*Bellezza et al., 2006*). The EPS from WT was clearly stained under both pH conditions, strongly suggestive of the sulfate modification.

### Gene cluster for the biosynthesis of viscous polysaccharides

*slr5054* resides on a megaplasmid, pSYSM, in a large gene cluster (*sll5042–60*), which we named *xss* (extracellular sulfated polysaccharide biosynthesis) (Fig. 2A, *xssA*–*xssS*). This cluster includes two genes for sulfotransferases (*xssA*, *xssE*), eight genes for glycosyltransferases (*xssB*, *xssC*, *xssG*, *xssI*, *xssM*, *xssN*, *xssO*, *xssP*), three genes for the polysaccharide polymerization system (Wzx/flippase; *xssH*, Wzy/polymerase; *xssF*, and polysaccharide co-polymerase [PCP]; *xssK*), one gene for a putative transcriptional regulator (*xssQ*), a pair of genes for the bacterial two-component phosphorelay system (*xssR*, *xssS*), and genes encoding several small proteins of unknown function (Table S1, Fig. 2A). All genes except those of unknown function were disrupted individually with a read-through cassette, and segregation was confirmed by colony PCR (Fig. S4). Bloom formation and sugar content of the EPS fraction were reduced in many mutants (Fig. 2B, C). In particular, bloom formation was completely abolished in Δ*xssA*, Δ*xssB*, Δ*xssF*, Δ*xssH*, Δ*xssK*, Δ*xssM*, Δ*xssN*, and Δ*xssP*, in which EPS accumulation was also suppressed. Certain glycosyltransferase mutants (Δ*xssC*, Δ*xssG*, Δ*xssI*, *ΔxssO*) formed blooms but accumulated little EPS, and neither bloom formation nor EPS accumulation was substantially altered in one sulfotransferase mutant (Δ*xssE*). In general, the Wzx/Wzy system in bacteria produces various EPS, lipopolysaccharides, and CPS through four steps: (i) biosynthesis of a heterooligosaccharide repeat unit on a lipid linker at the cytoplasmic side of the plasma membrane by a series of glycosyltransferases and modification enzymes, (ii) flip-out of the unit to the periplasmic side by Wzx, (iii) polymerization by transfer of the nascent polysaccharide chain to the repeat unit by Wzy, and (iv) export of the EPS chain through the periplasm and outer membrane via PCP and the outer-membrane polysaccharide export protein (OPX) (*Islam and Lam, 2014, Schmid et al., 2015*). It is very likely that the *xss* cluster harbors a whole set of genes for the Wzx/Wzy-dependent pathway except a gene for OPX.

**Figure 2.**
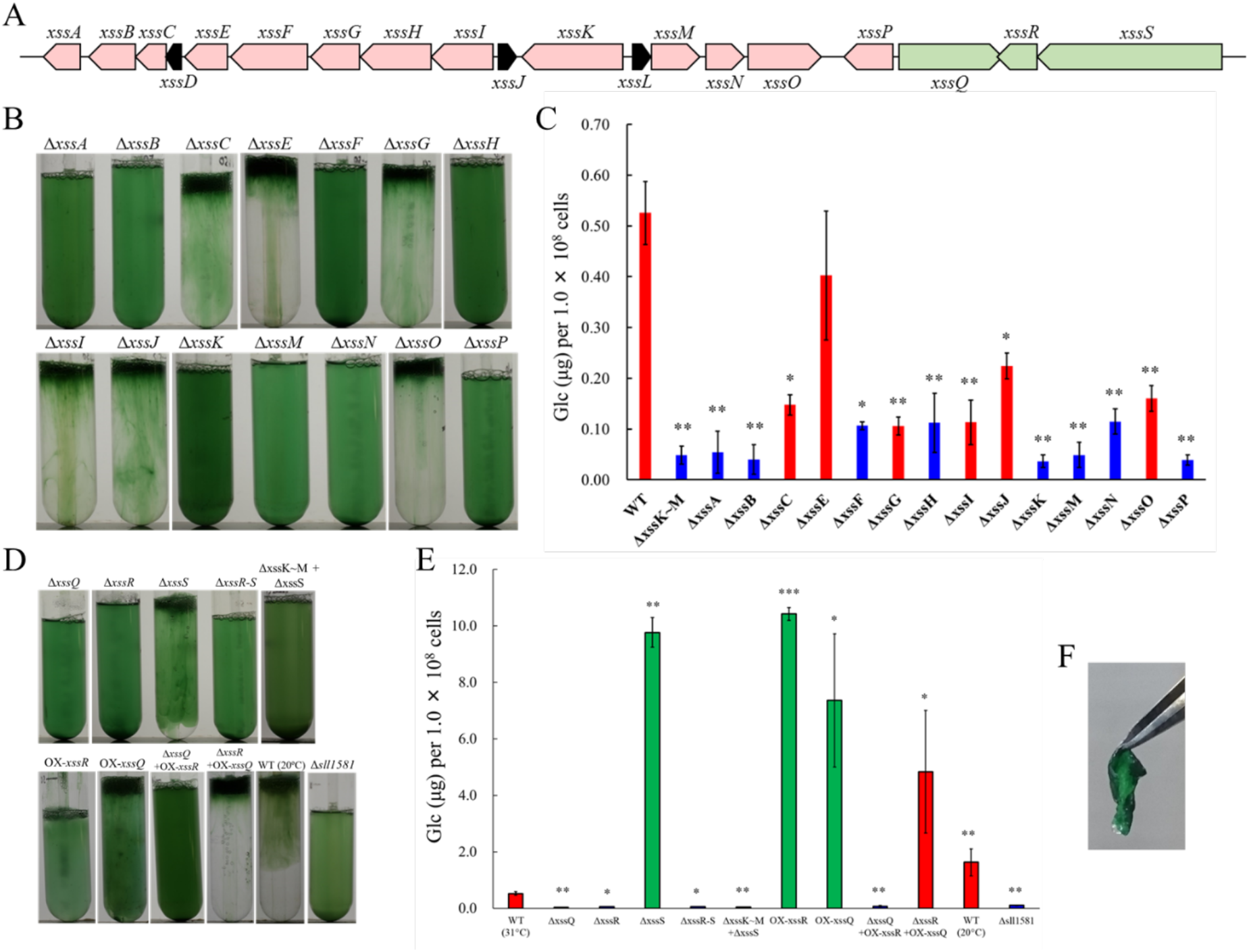
The *xss* gene cluster and phenotype of *xss* mutants. **A,** The *xss* gene cluster. Red, polysaccharide biosynthesis genes; green, regulatory genes; black, genes of unknown function. **B,** Bloom formation by the mutants carrying disruptions in the polysaccharide biosynthesis *xss* genes. **C**, Total sugar content (μg glucose per 1 × 10^8^ cells) of the EPS fraction from mutants in b. Red bars, bloom-forming mutants; blue bars, non-bloom-forming mutants. Error bars represent SD (WT grown at 20°C, n = 6; others, n = 3). Statistical significance was determined using Welch’s *t* test (\**P* < 0.05, \*\**P* < 0.005, \*\*\**P* < 0.0005). **D,** Bloom formation by regulatory mutants, WT grown at 20°C, and OPX mutant (Δ*sll1581*). **E,** Total sugar content of the EPS fraction from mutants in d. Red bars, bloom-forming mutants; green bars, excess-bloom-forming mutants. **F,** A sheet of OX-*xssR* cells was stripped off from the agar plate by tweezers. The culture temperature was 31°C unless otherwise stated.

### Regulation of the sulfated EPS biosynthesis

The sensory histidine kinase mutant Δ*xssS* accumulated a much larger amount of EPS than WT, whereas mutants of the cognate response regulator *xssR* and transcriptional regulator *xssQ* had a null phenotype with regard to both bloom formation and EPS accumulation (Fig. 2D, E, Table S2). The double mutant Δ*xssS*/Δ*xssR* had a phenotype similar to that of Δ*xssR*. Overexpression of *xssR* or *xssQ* (OX-*xssR*, OX-*xssQ*) resulted in strong bloom formation as well as marked accumulation of viscous EPS, similar to that seen for Δ*xssS*. The combination of *xssQ* disruption and *xssR* overexpression (Δ*xssQ+*OX-*xssR*) abrogated bloom formation and EPS accumulation, whereas the combination of *xssQ* overexpression and *xssR* disruption (Δ*xssR+*OX-*xssQ*) resulted in a pronounced phenotype of bloom formation and EPS accumulation. These results suggested that the sensor histidine kinase XssS suppresses the response regulator XssR, leading to activation of the transcriptional activator XssQ. Notably, the OX-*xssR* and OX-*xssQ* strains formed sticky, non-motile, biofilm-like colonies on agar plates that could be picked by tweezers (Fig. 2F).

XssQ is a new type of the <u>s</u>ignal <u>t</u>ransduction <u>A</u>TPase with numerous domains (STAND) protein, because it harbors an N-terminal helix-turn-helix transcriptional DNA-binding domain (Fig. S5). Typical STAND proteins possess a three-domain module with ATPase activity and are involved in processes such as apoptosis and immunity in animals, plants, and some bacteria (*Danot et al., 2009*). Using real-time quantitative PCR (qPCR), we compared gene expression in the *xss* cluster for WT, Δ*xssS*, and Δ*xssQ* (Fig. 3A). Expression of five genes (*xssA*, *xssB*, *xssE*, *xssN*, *xssP*) was very low in Δ*xssQ* compared with WT, whereas that of *xssF*, *xssH*, and *xssK* was not substantially affected. These results suggested that XssQ transcriptionally activates genes encoding sulfotransferases and certain glycosyltransferases but not genes for polymerization and export via the Wzx/Wzy system. qPCR analysis of gene expression in Δ*xssS* revealed a tendency for slight upregulation of *xssB*, *xssE*, *xssN*, and *xssP*. We performed RNA-seq analysis of WT, Δ*xssS*, and Δ*xssQ* to see the transcriptome (Fig. S6). The genes down-regulated in Δ*xssQ* and up-regulated in Δ*xssS* were mostly *xss* genes. In detail, the regulated genes were *xssA-E* and *xssL-P*, which were roughly consistent with the qPCR analysis. We conclude that *xssA-E* and *xssL-P* were specifically regulated by XssS/XssR/XssQ. In a previous report, *xssA*–*xssE* and *xssL*–*xssP* were up-regulated at low temperature in another substrain of *Synechocystis* 6803 (*Kopf et al., 2014b*). To test this in our substrain, we measured the sulfated EPS accumulation of WT culture at 20°C, and it was 3.1-fold greater than that at normal growth temperature (31°C; Fig. 2D, E and Table S2). This result suggests that XssS/XssR/XssQ is a unique temperature sensor for *xss* gene expression.

**Figure 3.**
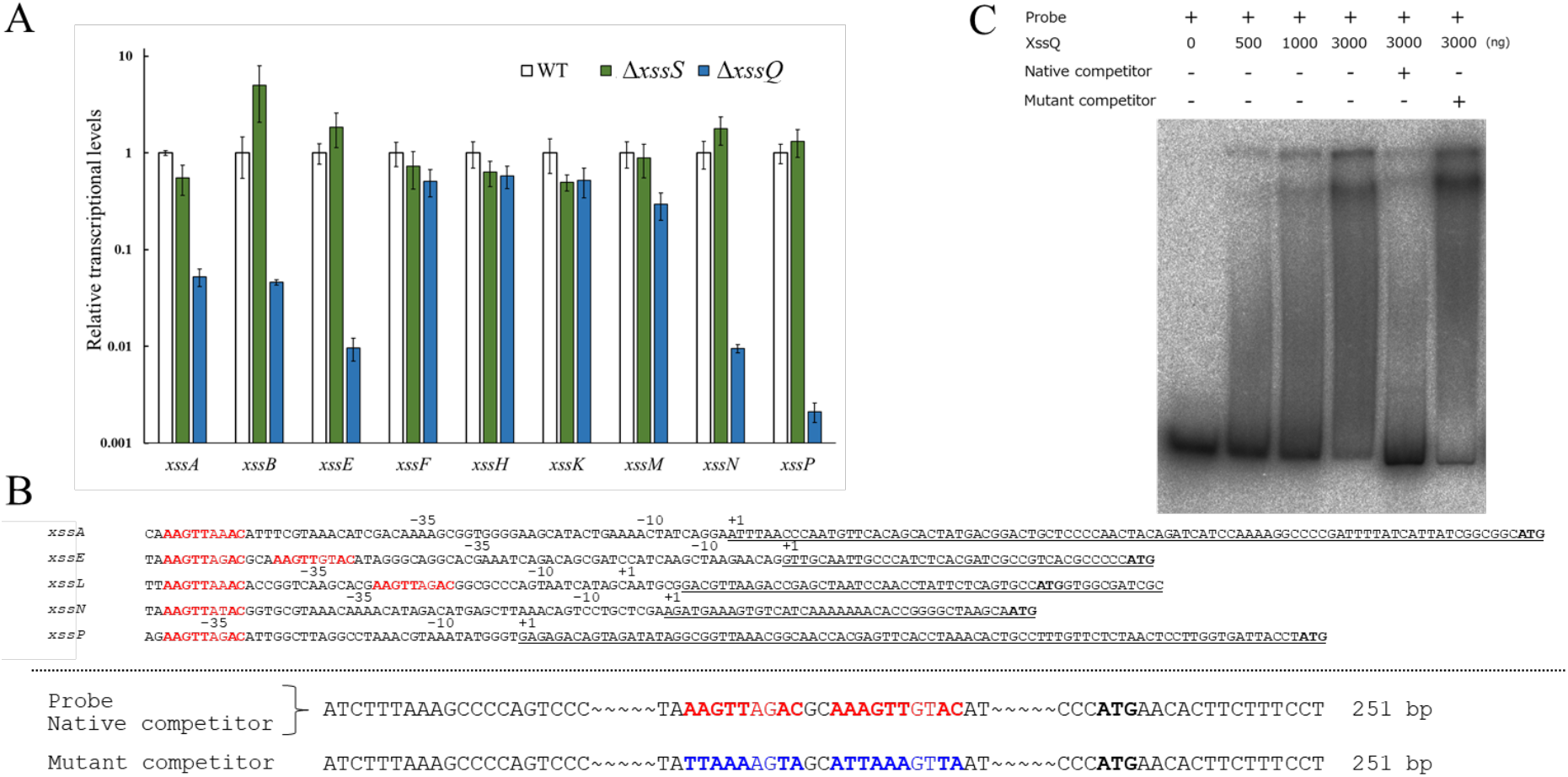
Transcriptional regulation of *xss* genes. **A,** Transcript levels for *xss* genes in WT, Δ*xssS*, and Δ*xssQ* measured by qPCR. The internal standard was *rnpB*. Relative expression levels were obtained by normalization to the transcript levels of each gene in WT. Error bars represent SD (n = 3, biological triplicates). **B**, upper; Sequence comparison of upstream regions of the five regulated genes. Putative consensus regions are shown in red and fully conserved nucleotides are shown in bold letter. Underlines represent transcribed regions based on the report(*Kopf et al., 2014b*) and initiation codons of regulated genes are shown in black/bold letters. **B**, lower: Sequences of DNA probe and competitors for *xssE* (native and mutant) used for EMSA of C. Consensus regions are shown in red, and mutated region are shown in blue. Total DNA size is 251 bp, where identical sequences are mostly not shown except 20 bp at both ends. **C,** The autoradiogram image of EMSA of XssQ protein and the DNA probe of *xssE* with some competitors.

We aligned nucleotide sequences near the transcription start sites of the regulated genes (*xssA, xssE, xssL, xssN,* and *xssP*) to find the consensus sequences for XssQ binding (Fig. 3B), according to the differential RNA-seq-type transcriptomic analysis of *Synechocystis* 6803 (*Kopf et al., 2014b*). There are single or tandem consensus sequence, AAGTTXXAC. To confirm the binding of XssQ to this region, we performed electrophoretic mobility shift assay (EMSA) using purified recombinant XssQ protein and a PCR-amplified DNA fragment of *xssE* upstream (Fig. 3C). The band position of the radiolabeled probe DNA shifted reflecting the concentration of XssQ. This shift was largely eliminated by excess addition of the unlabeled native competitor, but not by addition of the mutant competitor with mutations in the consensus region. These results suggest that XssQ recognizes the consensus sequence of *xssE* and other target genes for their transcriptional activation.

### The chemical composition of the sulfated EPS

EPS of WT and the 19-fold overproduction mutant (Δ*xssS*) were subjected to chemical composition analysis (Table 1, Fig. S7). EPS from WT included various sugars and some sulfate groups, whereas EPS from Δ*xssS* consisted of only four types of sugars and sulfate groups with the near stoichiometric molar ratio of rhamnose:mannose:galactose:glucose:sulfate of 1:1:1:5:2. This finding roughly fits with the gene number, i.e., eight glycosyltransferase genes and two sulfotransferase genes. We speculated that the overaccumulation of EPS in Δ*xssS* reflects the true product of the *xss* cluster. The EPS from WT may contain a considerable amount of unrelated polysaccharides, which were erroneously recovered together with the *xss* product. Here, the sulfated EPS produced by the *xss* cluster in *Synechocystis* 6803 was designated “Synechan”.

**Table 1.**
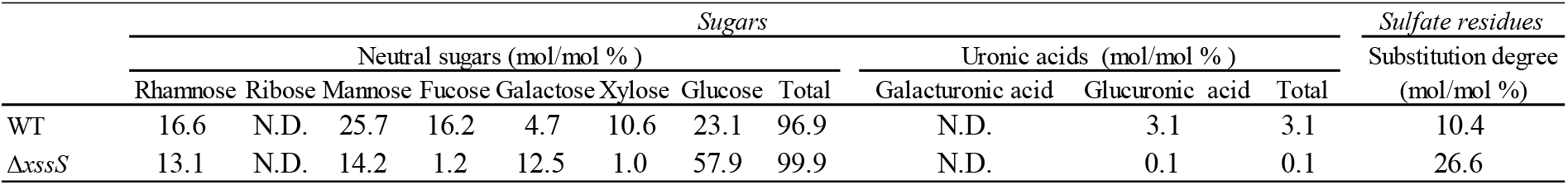
Chemical composition of the EPS from WT *Synechocystis* 6803 and Δ*xssS* mutant.

### The OPX protein for synechan biosynthesis

There is no candidate gene in the *xss*-carrying plasmid for the OPX protein of the Wzx/Wzy system, whereas *sll1581*, an OPX homolog, was found on the main chromosome. Disruption of *sll1581* (Δ*sll1581*) abolished bloom formation and EPS accumulation (Fig. 2D, E). Thus, the chromosomal OPX protein Sll1581 (XssT) appears to serve as the outer-membrane exporter for synechan. Interestingly, *Synechocystis* 6803 possesses *xssT* (OPX gene) and *sll0923* (a second PCP-2a gene) on the main chromosome and *xssK* (PCP-2a gene) on the plasmid pSYSM, whereas its close relative *Synechocystis* 6714 harbors only homologs of *sll0923* and *xssT* but lacks the entire plasmid carrying the *xss* cluster. This suggested that XssT serves as an OPX for dual function for both XssK and Sll0923. It is likely that *Synechocystis* 6803 acquired pSYSM and borrowed the chromosomal OPX gene *xssT* to produce synechan or, alternatively, *Synechocystis* 6714 may have lost the plasmid.

## Discussion

Summarizing these data, we propose models for synechan biosynthesis apparatus including OPX and temperature-responsive regulation (Fig. 4A, B and Fig. S5). The model of the Xss apparatus fits well with the known Wzx/Wzy-dependent apparatus represented by xanthan biosynthesis in *Xanthomonas campestris* (*Katzen et al., 1998*). The eight glycosyltransferases including XssP (the priming glycosyltransferase) produce oligosaccharide repeat unit of eight sugars, which is consistent with the sugar composition of synechan. These findings suggest that the *xss* cluster on the pSYSM plasmid harbors a whole set of genes for synechan biosynthesis except the OPX gene (*xssT* on the main chromosome). Notably, the cluster harbors two sulfotransferase genes, which have not been found to our knowledge in other bacterial gene clusters for extracellular polysaccharide biosynthesis. Sulfotransferases, XssA and XssE, belong to distinct subfamilies of bacterial sulfotransferases. We found many sulfotransferase genes in various cyanobacterial genomes by Pfam search (PF00685, PF03567, PF13469). They are mostly found in gene clusters for putative extracellular polysaccharide biosynthesis (Wzx/Wzy-type and ABC-type) (Fig. S8). It should be noted that they are more or less partial as a cluster for extracellular polysaccharide biosynthesis system, whereas the *xss* cluster appears to be complete except the OPX gene in *Synechocystis* 6803. It is well established that the polysaccharide moiety of membrane-anchored lipopolysaccharides and CPS of bacteria are produced and exported by the Wzx/Wzy-dependent or ABC transporter–dependent pathways, whereas free EPS, i.e., xanthan and cellulose, are produced by the Wzx/Wzy-dependent and synthase-dependent pathways but not by the ABC transporter–dependent pathway (*Schmid et al., 2015, Willis and Whitfield, 2013*). In the literature, a sulfated CPS was reported in *Arthrospira platensis* (formerly *Spirulina platensis*) (*Mouhim et al., 1993*). This sulfated CPS may be produced by an ABC transporter-type gene cluster in Fig. S8. Gene disruption will confirm such predictions deduced from the gene cluster analyses, although targeted disruption is not so easy in many cyanobacteria due to poor transformation efficiency except *Synechocystis* 6803. In contrast, no sulfated polysaccharide has been reported in the other bacteria, though many sulfotransferases are also registered in Pfam database. Some of them are known to transfer a sulfuryl group to lipo-oligosaccharides in rhizobia (Nod factor) and mycobacteria (*Mougous et al., 2002*).

**Figure 4.**
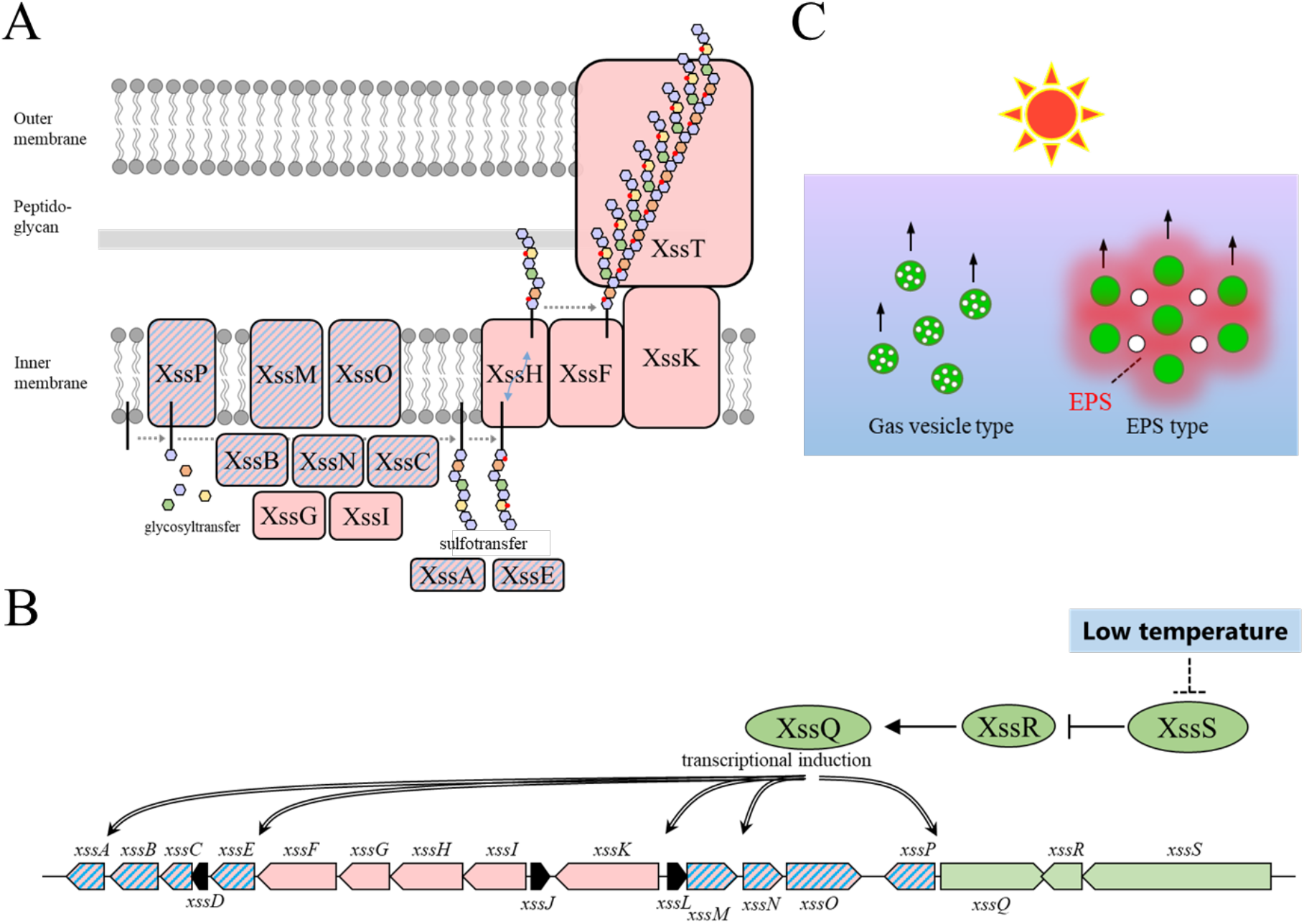
Proposed models for the synechan biosynthesis apparatus, transcriptional regulation, and bloom formation. **A,** Model for the synechan biosynthesis apparatus with sugar polymerization and modification. Red boxes represent biosynthesis components, and red boxes with blue stripes represent transcriptionally regulated components. A putative lipid linker is shown as a black rod. Each monosaccharide is shown as a small hexagon, and each sulfate group is shown as a red spot. **B,** Signaling and transcriptional regulation model. Green arrows and ellipses represent regulatory genes and proteins, respectively. Genes for synechan biosynthesis are shown in red, and transcriptionally regulated genes are depicted with blue stripes. Arrows with double lines represent transcriptional activation. **C,** Two flotation models for bloom formation in cyanobacteria. Left, flotation of cells (green circles) with intracellular gas-filled vesicles (white circles). Right, flotation of EPS (red shading)-entrapped cells (green circles) and extracellular gas bubbles (white circles), which are generated by photosynthesis.

XssQ, a STAND protein with a DNA binding domain is indeed the transcriptional activator for *xssE* and other induced genes. XssQ homologs are found widely throughout the cyanobacteria but the set of XssS/XssR/XssQ is found near the gene cluster for sulfated EPS biosynthesis with sulfotransferases in many cyanobacteria (Fig. S9, Table S3). Consensus sequences are also found in upstream of some genes in the cluster, suggesting that the XssS/XssR/XssQ system may operate universally for induction of sulfated EPS production under certain environmental conditions such as cold temperature.

Acidic polysaccharides containing uronic acids and other carboxylic groups are common in bacteria, but sulfated polysaccharides are produced exclusively by cyanobacteria (*Pereira et al., 2009*). To speculate on the physiological significance of sulfated polysaccharides, we summarized the distribution of sulfotransferase genes in cyanobacteria of various habitats (Table S3). Species living in high salinity environments such as a salt lake or seawater mostly produce sulfated polysaccharides. The freshwater species *Synechocystis* sp. PCC 6714 lacks the *xss* genes including sulfotransferases and salt-resistance genes that are present in the more salt-resistant *Synechocystis* 6803 (*Kopf et al., 2014a*). A cyanobacterial sulfated polysaccharide, sacran, shows much higher capacity for saline absorption than does hyaluronic acid, a uronic acid–containing polysaccharide, whereas both absorb pure water efficiently (*Okajima et al., 2008*). The wide distribution of sulfated polysaccharides among many cyanobacteria may reflect their inherent compatibility with survival in some saline environments.

There are many putative genes that may be involved in biosynthesis and export of extracellular polysaccharides and lipopolysaccharides in the genome of *Synechocystis* 6803 (*Fisher et al., 2013, Pereira et al., 2015*). Many of them have been disrupted for characterization. With regard to the Wzx/Wzy-dependent system, *sll5052* (*xssK*) disruption mutant showed no clear phenotypes for bloom formation or EPS production (*Jittawuttipoka et al., 2013*), probably because the parent strain did not produce discernable amount of EPS like our nonmotile strain. Similarly, the deletion mutant of *sll5049* (*xssH*) did not show any defect in EPS or CPS accumulation, though related mutants (Δ*sll0923* for a second PCP-2a) were shown to be depleted slightly of both CPS and EPS (*Pereira et al., 2019*). These results contrast with our null phenotype of Δ*sll5052* (*xssK*) and Δ*sll5049* (*xssH*), probably because of the difference in the parent strains. In addition, we found that our Δ*sll0923* did not show any defect in the bloom formation. On the other hand, disruption of *sigF* (*slr1564*) for a sigma factor of global cell surface regulation increased three to four fold accumulation of sulfated EPS (*Flores et al., 2019a, Flores et al., 2019b*). The proteome analysis of Δ*sigF* revealed many (more than 160) proteins except for any Xss proteins were up-regulated, leaving the sulfated EPS biosynthesis pathway elusive. The sugar composition of the sulfated EPS of their WT is similar to our WT, although the composition of EPS of Δ*sigF* was different for WT or synechan from our Δ*xssS*.

To get insights into the difference in bloom formation between the motile and nonmotile substrains, we compared transcription data (Table S4). It is evident that many *xss* genes on the plasmid are expressed several times higher in the motile substrain than the nonmotile one except for *xssT* on the main chromosome, despite that the nucleotide sequence of the *xss* gene cluster was completely conserved between them. This fact suggests a possibility that another mechanism besides the XssS/XssR/XssQ contributes to the difference between the substrains. For example, the plasmid copy number of pSYSM may be higher in the motile substrain than the nonmotile substrain. The plasmid function is often affected depending on variations in the main chromosome (*Vial and Hommais, 2020*). Anyway, it is very important to select the parent strain depending on the research purpose.

The cyanobacterial bloom rapidly accumulates in populations of cyanobacterial cells floating on the water surface, which often produce potent cyanotoxins (hepatotoxins, neurotoxins, etc.) (*Merel et al., 2013*). Blooms are thought to be supported mainly by cellular buoyancy due to intracellular proteinaceous gas vesicles constructed by gas vesicle proteins (*Beard et al., 2002, Walsby, 1994*). Moreover, recent studies suggested that extracellular polysaccharides are also important for the bloom formation (*Chen et al., 2019*). Some papers reported that the cells without gas vesicle can form blooms by EPS-dependent manner after artificial addition of divalent cations (Ca^2+^ or Mg^2+^) (*Dervaux et al., 2015, Wei et al., 2019*). On the other hand, our study demonstrated that the gas-trapped EPS is sufficient for bloom formation of *Synechocystis* 6803, which does not produce gas vesicles, without addition of any divalent cations. This result is consistent with reports that cyanobacteria without gas vesicles form booms in natural environments, including freshwater lakes (*Casero et al., 2019, du Plooy et al., 2015, Steffen et al., 2012*).

Finally, sulfated polysaccharides are expected to be healthy foods, industrial materials and medicines (*Jiao et al., 2011, Wardrop and Keeling, 2008*). Some sulfated EPS from *Synechocystis* 6803 showed antitumor activity (*Flores et al., 2019a*), though this EPS may not be identical to synechan. The Xss-dependent biosynthesis of synechan in *Synechocystis* 6803 should be a good model for studies of other cyanobacterial sulfated polysaccharides. Combinatorial expression of sulfotransferases and glycosyltransferases from other cyanobacteria in *Synechocystis* cells will provide clues to their functions. Heterologous expression of *Synechocystis xss* genes in other organisms will also open a possibility of large scale production of modified synechan species. Further molecular studies of *xss* genes and related genes from the database should accelerate screening and potential applications of cyanobacterial sulfated polysaccharides.

## Materials and Methods

### Cyanobacterial strains and cultures

The motile substrain PCC-P of the unicellular cyanobacterium *Synechocystis* sp. PCC 6803, which exhibits phototaxis (*Yoshihara et al., 2000*) and forms bloom-like aggregates, was used as the WT in this work. A non-motile glucose-tolerant substrain, which has been widely used for studies of photosynthesis, was used for comparison (*Chin et al., 2018*). Cells were maintained in BG11 liquid medium (*Stanier et al., 1971*) under continuous illumination with bubbling of 1% CO2 in air at 31°C, or on 1.5% agar plates. White light of 30 μmol photons m^−2^s^−1^ was generated by fluorescent lamps. Cell density was monitored at 730 nm.

### Construction of plasmids and mutants

Primers used are listed in Table S5. Plasmids and mutants were constructed as described (*Chin et al., 2018*). In brief, the DNA fragments, antibiotic-resistance cassettes, the *trc* promoter, and plasmid vectors were amplified by PCR using PrimeSTAR MAX DNA polymerase (Takara, Shiga, Japan) and combined using the In-Fusion System (Takara). The resulting plasmid constructs were confirmed by DNA sequencing.

Gene disruption was performed in two different ways. One method was replacement of a large portion of a targeted gene(s) with an antibiotic resistance cassette. The other method was replacement of the translation initiation codon with a stop codon. In both cases, the screening cassette without the terminator was inserted in the direction of the targeted gene(s) to allow transcriptional readthrough of the downstream gene(s). For overexpression, gene expression was constitutively driven by the strong *trc* promoter in two ways: integration of a target gene with the strong *trc* promoter into a neutral site near *slr0846* or IS203c, or replacement of the target-gene promoter with the *trc* promoter. Natural transformation and subsequent homologous recombination were performed as described(*Chin et al., 2018*). The antibiotic concentration for the selection of transformants was 20 μg·mL^−1^ chloramphenicol, 20 μg·mL^−1^ kanamycin, and/or 20 μg·mL^−1^ spectinomycin. Complete segregation of the transformed DNA in the multicopy genome was confirmed by PCR using primers listed in Table S5, and the transformants are listed in Table S6.

### Bloom formation

The bloom was reproducibly formed using the two-step culture regime we developed in this work. Before the bloom formation experiment, cells were precultured once in liquid after transfer from plates. In the first step, cells inoculated at OD_730_ = 0.2 were grown with vigorous aeration under continuous light at 31°C or 20°C for 48 h. Typically the cell density reached OD_730_ ~2. In the second step, the culture was shifted to the standing condition without bubbling under the same continuous light for another 48 h (or longer) for cells to rise to the surface. Regarding the mutants of transmembrane glycosyltransferases, bloom formation was examined after 168 h of the second-step culture. The final concentration of the photosynthesis inhibitor DCMU (3-(3,4-<u>d</u>i<u>c</u>hlorophenyl)-1,1-di<u>m</u>ethyl<u>u</u>rea) was 100 μM.

### EPS fractionation

The fractionation method to isolate the crude EPS is shown in Fig. S1A. The viscous materials including cells after the second step of culture were collected by filtration using a 1.0-μm pore PTFE membrane (Millipore). The trapped materials were gently and carefully recovered from the membrane using MilliQ water with the aid of flat-tip tweezers. The collected sample was vortexed and then centrifuged at 20,000 × *g* for 10 min to remove cells. The supernatant constituted the crude EPS that contained viscous EPS and possibly CPS.

The refined fractionation method to isolate EPS is shown in Fig. 1F. The entire culture at the end of the first step, which did not contain gas bubbles, was first centrifuged at 10,000 × *g* for 10 min to remove cells and CPS and then filtered through a 1.0-μm pore PTFE membrane. The trapped EPS was carefully recovered as described above. The flowthrough of the filtration was regarded as free polysaccharides, which were recovered by ethanol precipitation. CPS was released from the cell pellet by vigorous vortexing with MilliQ water and recovered by centrifugation to remove cells (20,000 × *g* for 10 min).

### Sugar quantification

Total sugar was quantified using the phenol-sulfate method (*DuBois et al., 1956*). A 100-μL aliquot of 5% (w/w) phenol was added to 100 μL of a sample in a glass tube and vortexed three times for 10 s. Then, 500 μL of concentrated sulfuric acid was added, and the tube was immediately vortexed three times for 10 s and then kept at 30°C for 30 min in a water bath. Sugar content was measured by absorption at 487 nm using a UV-2600PC spectrophotometer (Shimadzu, Japan, Tokyo). Any contamination of the BG11 medium was evident by slight background coloration. This background was subtracted on the basis of the extrapolation of absorption at 430 nm, where the coloration due to sugars was minimal. Glucose was used as the standard. Some EPS samples were highly viscous, so we vortexed and sonicated them before measurement. Statistical significance was determined using Welch’s *t* test.

### Sugar composition analysis

The collected EPS samples were dialyzed with MilliQ water and then freeze-dried for 3 days. Sugar composition was analysed by Toray Research Center, Inc. (Tokyo, Japan). A part of the fluffy sample (WT, 0.298 mg; Δ*xssS*, 0.203 mg) was dissolved in 200 μL of 2 M trifluoroacetic acid and hydrolysed at 100°C for 6 h. The treated sample was vacuum-dried with a centrifugal evaporator, redissolved in 400 μL MilliQ water, and filtered through a 0.22-μm pore filter. This sample was used for the analysis.

Monosaccharide composition was determined by HPLC with the LC-20A system (Shimadzu). For neutral sugars, the column was TSK-gel Sugar AXG (TOSOH, Japan) and the temperature was 70°C. The mobile phase was 0.5 M potassium borate (pH 8.7) at 0.4 mL/min. Post-column labelling was performed using 1% (w/v) arginine and 3% (w/v) boric acid at 0.5 mL/min, 150°C. For uronic acids, the column was a Shimpack ISA-07 (Shimadzu) and the temperature was 70°C. The mobile phase was 1.0 M potassium borate (pH 8.7) at 0.8 mL/min. Post-column labelling was performed using 1% (w/v) arginine and 3% (w/v) boric acid at 0.8 mL/min, 150°C. The detector was a RF-10AXL (Shimadzu), with excitation at 320 nm and emission at 430 nm. The standard curves were prepared for each monosaccharide with standard samples.

The SO_4_^2−^ content was determined by anion exchange column chromatography using the ISC-2100 system (Thermo Fisher Scientific, USA, Massachusetts). The column was eluted via a gradient of 0–1.0 M KOH. The separation column was IonPac ASI l-HC-4 μm (Thermo Fisher Scientific). Electric conductivity was used for detection.

### Alcian blue staining

The polysaccharides were stained with 1% Alcian blue 8GX (Merck) for 10 min in 3 % acetic acid (pH 2.5) or 0.5 N HCl (pH 0.5) as previously described (*Di Pippo et al., 2013*).

### qPCR

The qPCR was performed as described in our previous work (*Maeda et al., 2018*). Cells were harvested by centrifugation at 5000 × *g* for 10 min at 4°C. Cell disruption and RNA extraction were done using an RNeasy Mini kit for bacteria (Qiagen, Venlo, Netherlands). In addition, cells were disrupted five times by mechanical homogenization with zirconia beads (0.1-mm diameter) in a microhomogenizing system (Micro Smash MS-100, TOMY SEIKO, Tokyo, Japan) at 5,000 rpm for 40 s. For cDNA preparation, RNA was reverse-transcribed using random primers (PrimeScript RT reagent kit with gDNA eraser, Takara). Real-time PCR was performed using the THUNDERBIRD SYBR qPCR Mix (Toyobo) and the Thermal Cycler Dice Real Time System II (Takara). The transcript level in each strain was normalized to the internal control (*rnpB*). The primers used are listed in Table S5.

### *EMSA* ( electrophoretic mobility shift assay)

The expression and purification of recombinant His-tagged proteins and EMSA were performed as described in our previous works (*Hirose et al., 2010, Maeda et al., 2014*). In brief, His-tagged XssQ was expressed using pET28a vector system and *E. coli* C41(DE3) strain. The protein was purified by Histrap HP column (Cytiva, Tokyo, Japan) and AKTA prime system (Cytiva). For probe and native competitor, the upstream region of *xssE* was amplified with the primer set xssEup-1F/2R (total 251 bp). As a mutant competitor, the same region of the chemically synthesized DNA fragment containing mutations in the two consensus sequences was used for amplification with the same primer set as mention above. Labelling of the DNA probe, electrophoresis, and autoradiography were performed as described (*Midorikawa et al., 2009*). We incubated the aliquots of the XssQ protein (0, 500, 1000, or 3000 ng/lane) with the radiolabeled probe for 30 min at room temperature. For competition, 3000 ng of XssQ was incubated with the probe and 20 pmol of unlabeled competitors (native or mutant).

### RNA-seq analysis

RNA-seq analysis was performed as described in our previous work (*Ohbayashi et al., 2016*). Total RNA of *Synechocystis* 6803 was extracted as described in the qPCR protocol. The contaminated genome DNA was removed by TURBO DNA-free™ Kit (Thermo Fisher Scientific).

### Bioinformatics analysis

The sequences of the proteins were obtained from NCBI (http://www.ncbi.nlm.nih.gov/) and Pfam (http://pfam.xfam.org/) (*Finn et al., 2016*). The domain architecture was searched using the Simple Modular Architecture Research Tool, SMART (*Letunic et al., 2015*). Glycosyltransferase classifications were based on the CAZy database (http://www.cazy.org/) (*Henrissat, 1991, Lombard et al., 2013*). Amino acid sequence similarity was evaluated by NCBI BLAST search.

## Acknowledgements

This study was supported by Grants-in-Aid for JSPS Fellows 15J07605 and 19J01251 (to K.M.), the Japan Society for the Promotion of Science for Scientific Research (16H06558 to M.I.), and the JST for CREST program (JPMJCR1653 to M.I.).

---

**Figure S1.**
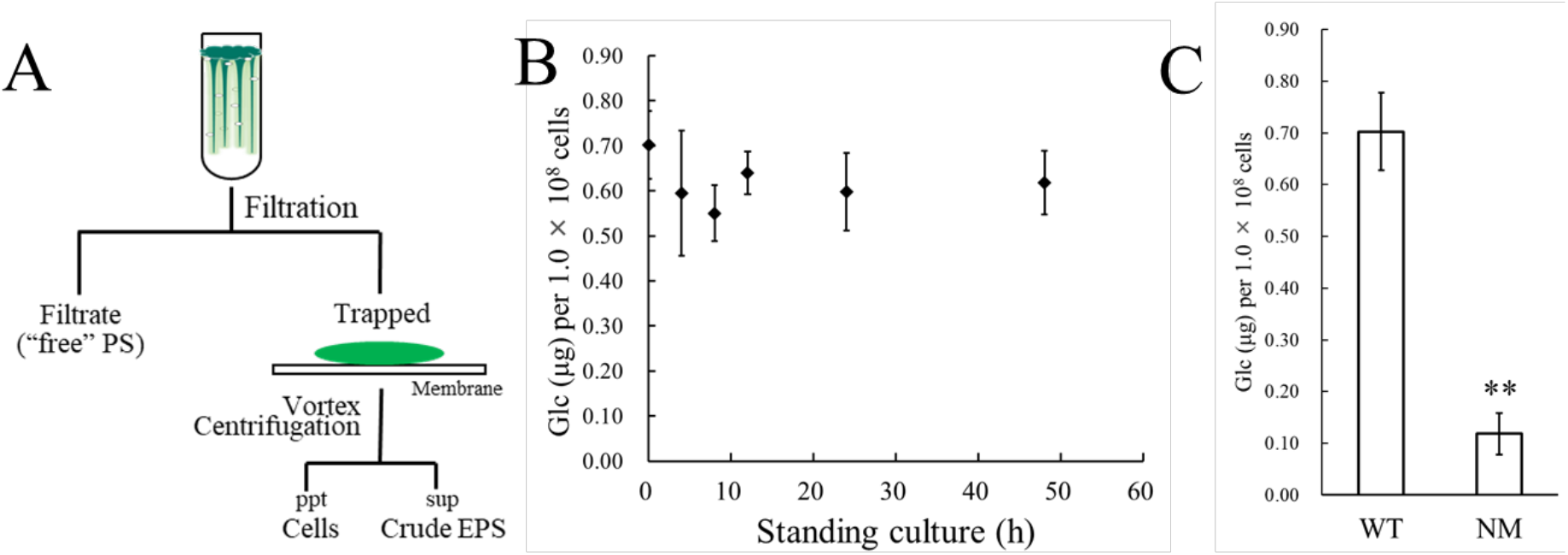
Isolation of crude EPS and sugar analysis. **A,** Protocol for isolating crude EPS from the bloomed culture. The bloom, including EPS and cells at the end of the second (standing) step of culture, was trapped by membrane filtration, and crude EPS was recovered from the bloom by vortexing and centrifugation. PS, polysaccharide; ppt, precipitate; sup, supernatant. **B,** Time course of sugar accumulation in the crude EPS during the standing culture. **C,** Sugar content of the crude EPS from WT, and a non-motile glucose-tolerant substrain (NM). Error bars represent SD (n = 3, \*\**P* < 0.005).

**Figure S2.**
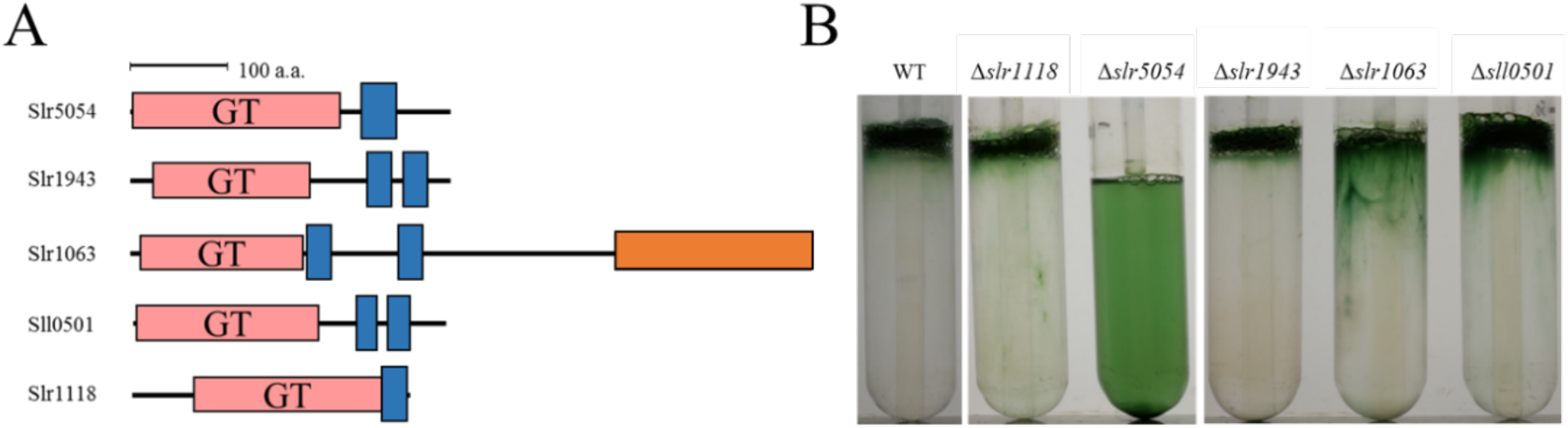
Bloom formation by several glycosyltransferase mutants. **A,** Domain architecture of membrane-bound glycosyltransferases. Red box, glycosyltransferase domain (GT); blue box, transmembrane region; orange box, glycogen phosphorylase domain. **B,** Bloom formation by mutants after the second step of culture.

**Figure S3.**
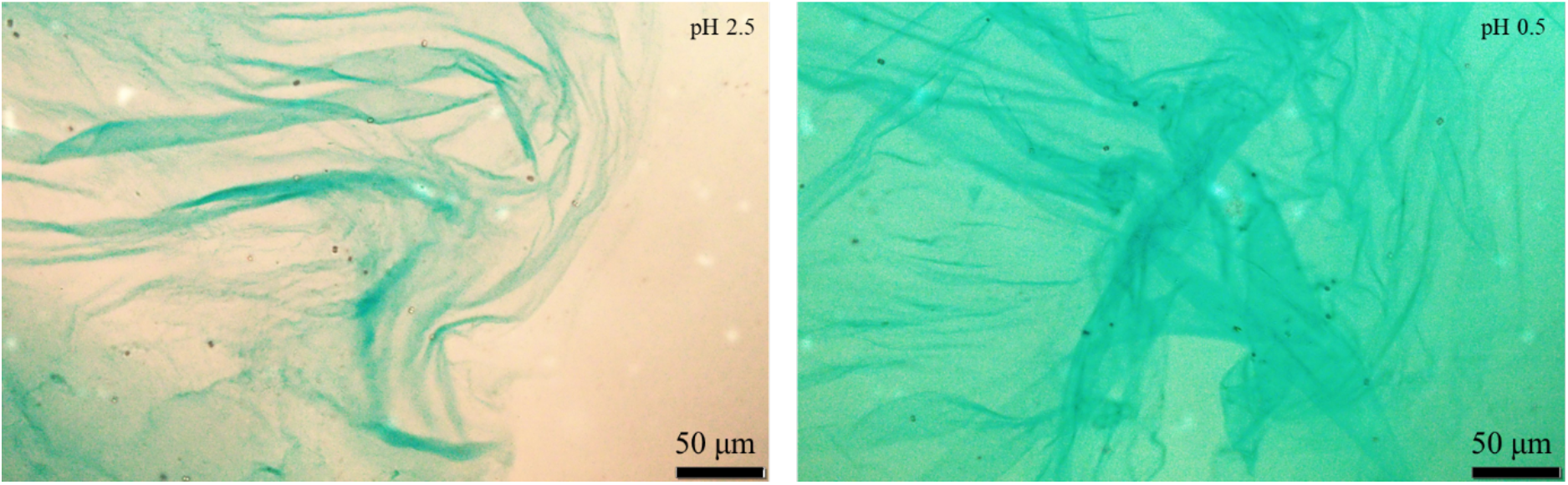
Alcian blue staining of EPS from WT. The microscopy images of isolated EPS from WT culture (Fig. 1g) stained with alcian blue at pH 2.5 (left) and pH 0.5 (right).

**Figure S4.**
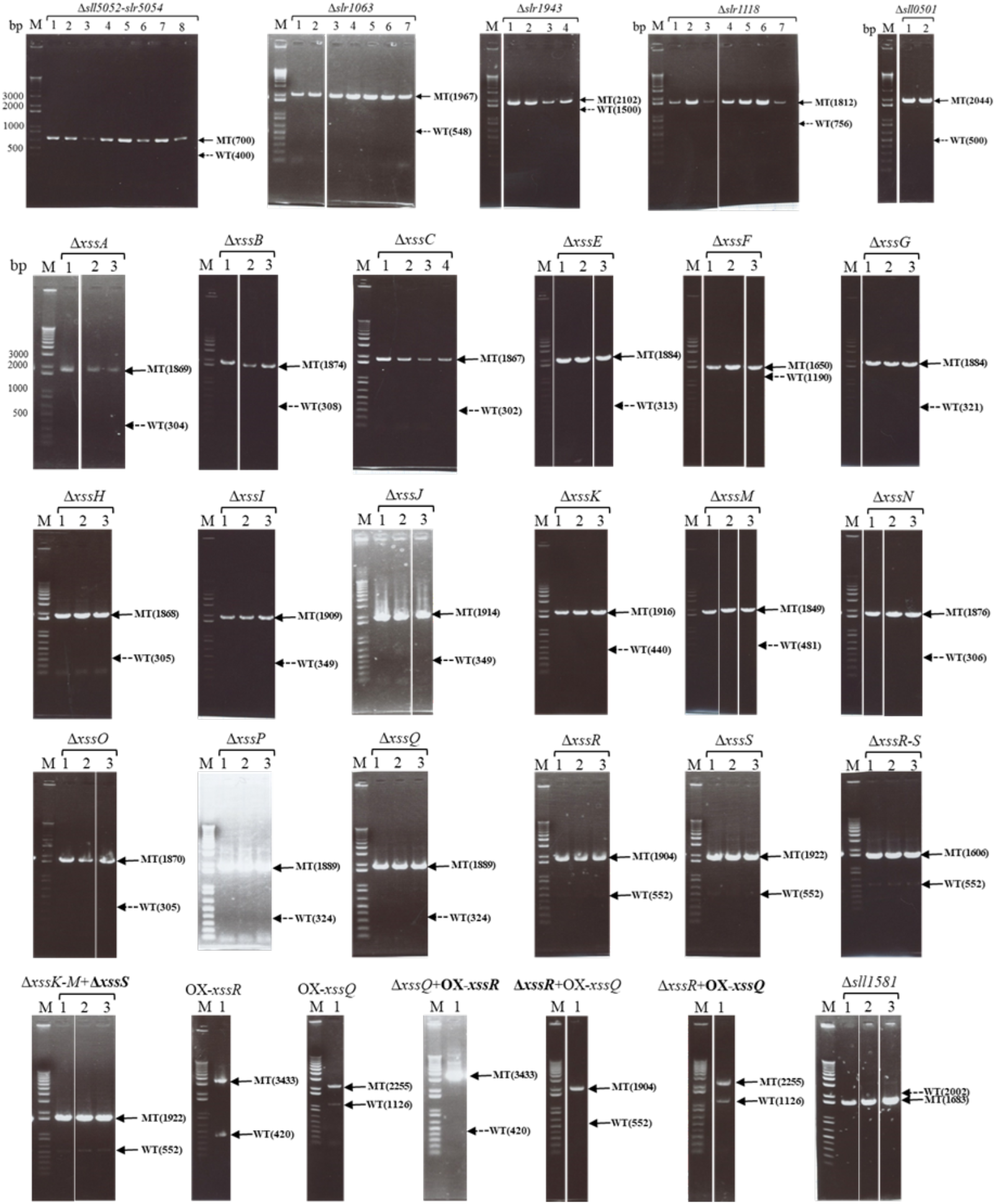
Agarose gel electrophoresis to assess segregation of mutants based on PCR data. M indicates marker lane, and the numbers above each lane indicate the different clones. The band positions of wild type (WT) and mutants (MT) are shown at the right, with theoretical lengths in nt. A solid arrow indicates the existence of the band; a dotted arrow indicates the absence of the band. The bold text in the strain names indicates the region assessed by PCR.

**Figure S5.**
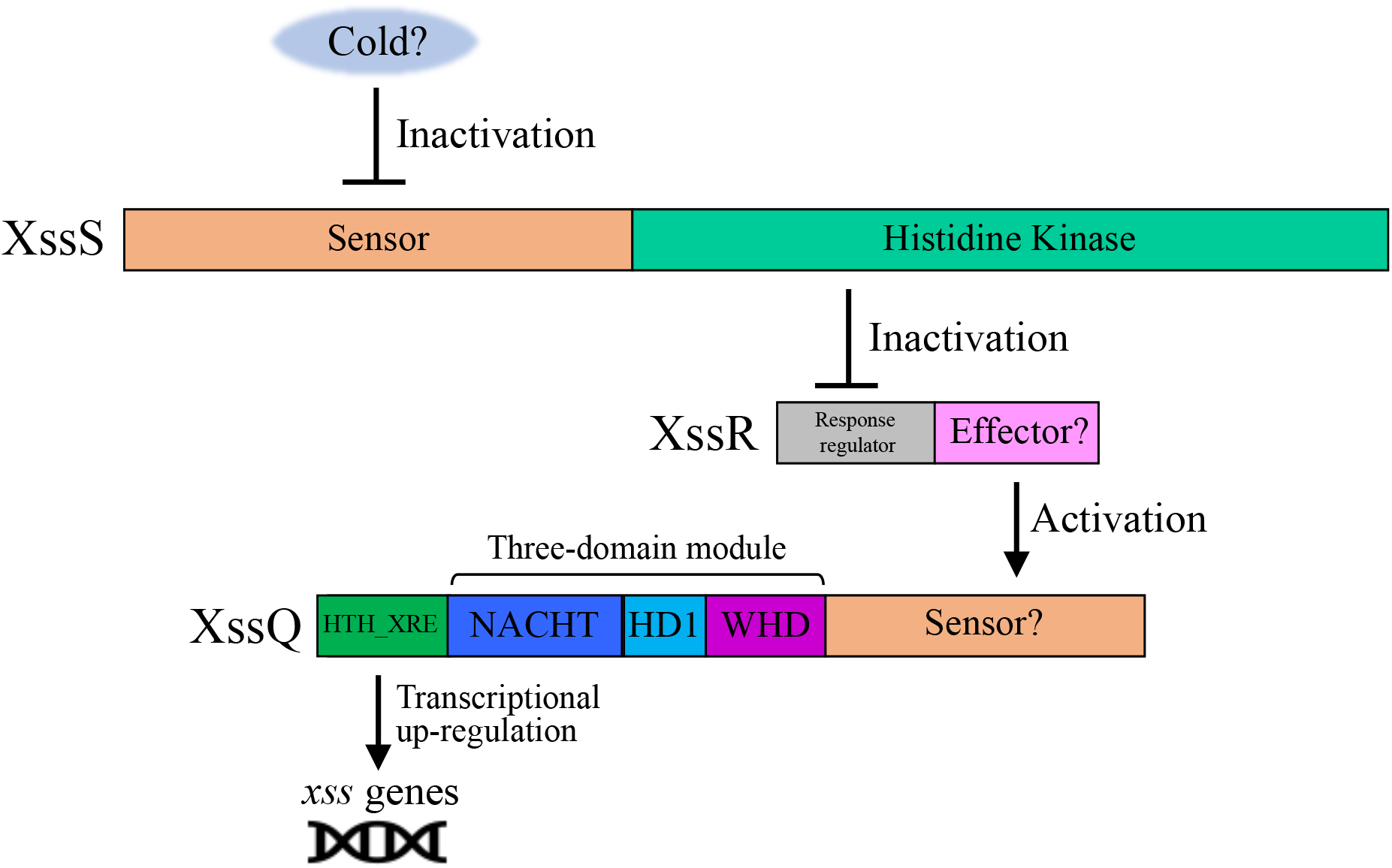
Domain architecture and proposed signal transduction pathway for XssS/XssR/XssQ. Hybrid-type histidine kinase XssS presumably senses a cold signal and transduces it to the response regulator XssR, which in turn activates the transcription of *xss* genes via the STAND protein XssQ. Generally, STAND proteins consist of the three-domain module, sensor region, and effector region^13^. In XssQ, the three-domain module consisting of the NACHT (<u>N</u>AIP [<u>n</u>euronal <u>a</u>poptosis <u>i</u>nhibitor <u>p</u>rotein], <u>C</u>IIA [MHC <u>c</u>lass <u>II</u> transcription <u>a</u>ctivator], <u>H</u>ET-E, and <u>T</u>P1 [mammalian <u>t</u>elomerase-associated proteins]) domain (PF05729), HD1 (<u>h</u>elical <u>d</u>omain <u>1</u>), and WHD (<u>w</u>inged <u>h</u>elix <u>d</u>omain) responds to a signal from XssR and oligomerizes, leading to the activation of the N-terminal effector domain (helix-turn-helix XRE family domain [SM00530]).

**Figure S6.**
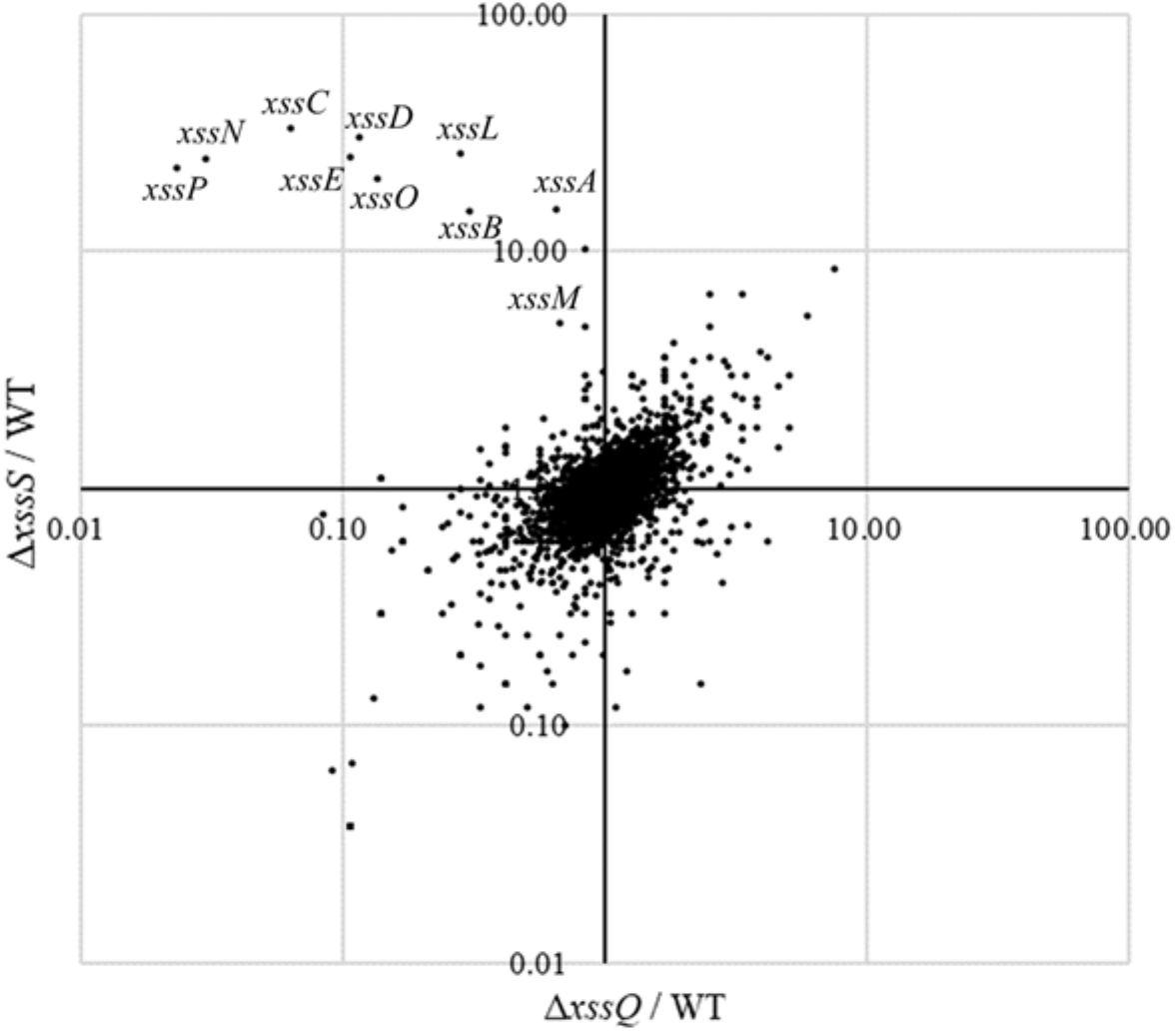
Scatter plot of transcriptome comparison among WT, Δ*xssS*, and Δ*xssQ*. Transcriptional levels were based on RPKM (Reads Per Kilobase of exon per Million mapped reads) value shown in supplementally data 1.

**Figure S7.**
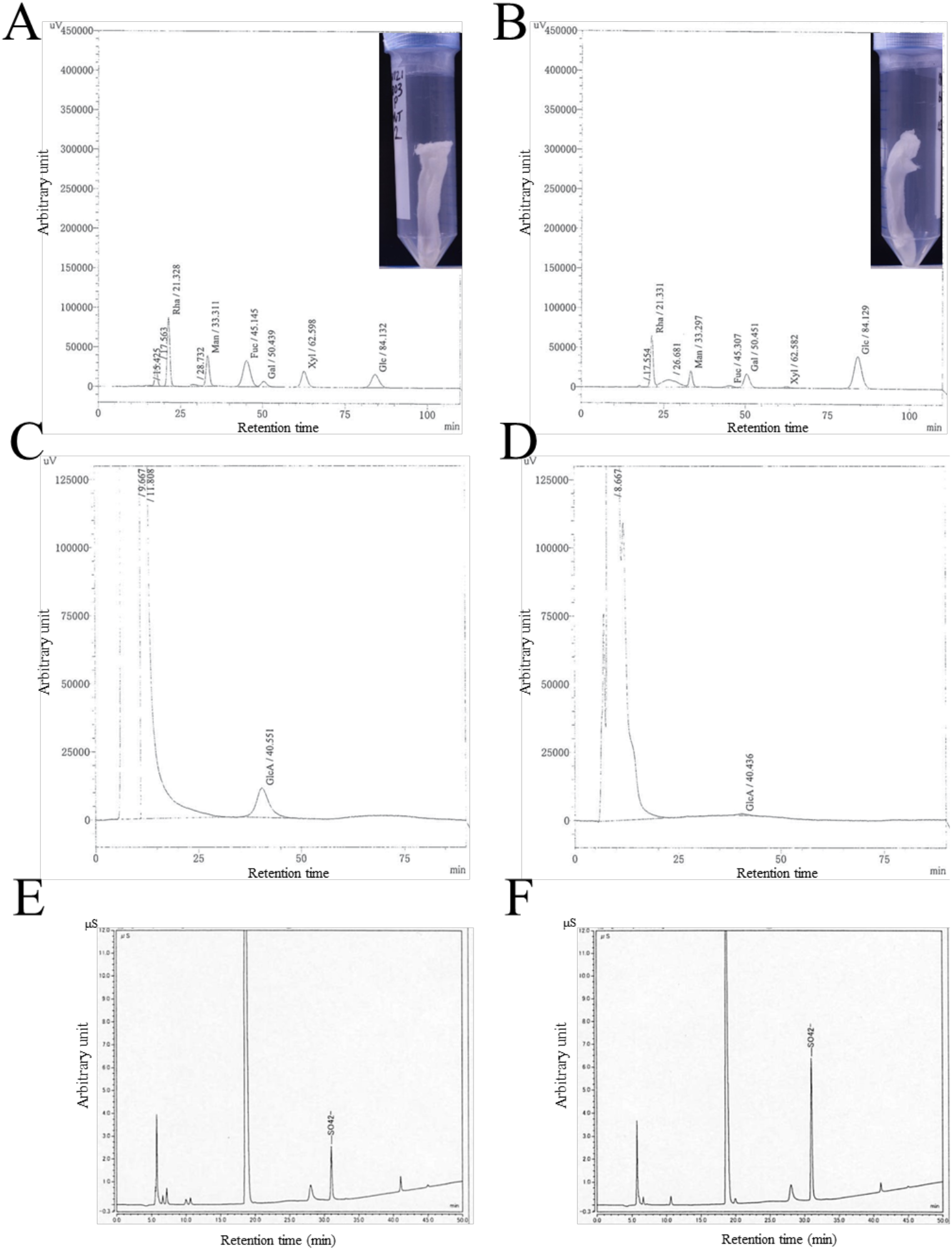
Chromatograms of HPLC and anion exchange column chromatography of EPS. **A** and **B**, HPLC profiles for neutral sugars in wild-type EPS (A) and Δ*xssS* EPS (B). **c** and **d**, HPLC profiles for uronic acids in wild-type EPS (C) and Δ*xssS* EPS (D). The corresponding monosaccharide and retention time are noted at each peak. **E** and **F**, HPLC profiles for SO4^2–^ after hydrolysis of EPS samples of wild-type (E) and Δ*xssS* (F).

**Figure S8.**
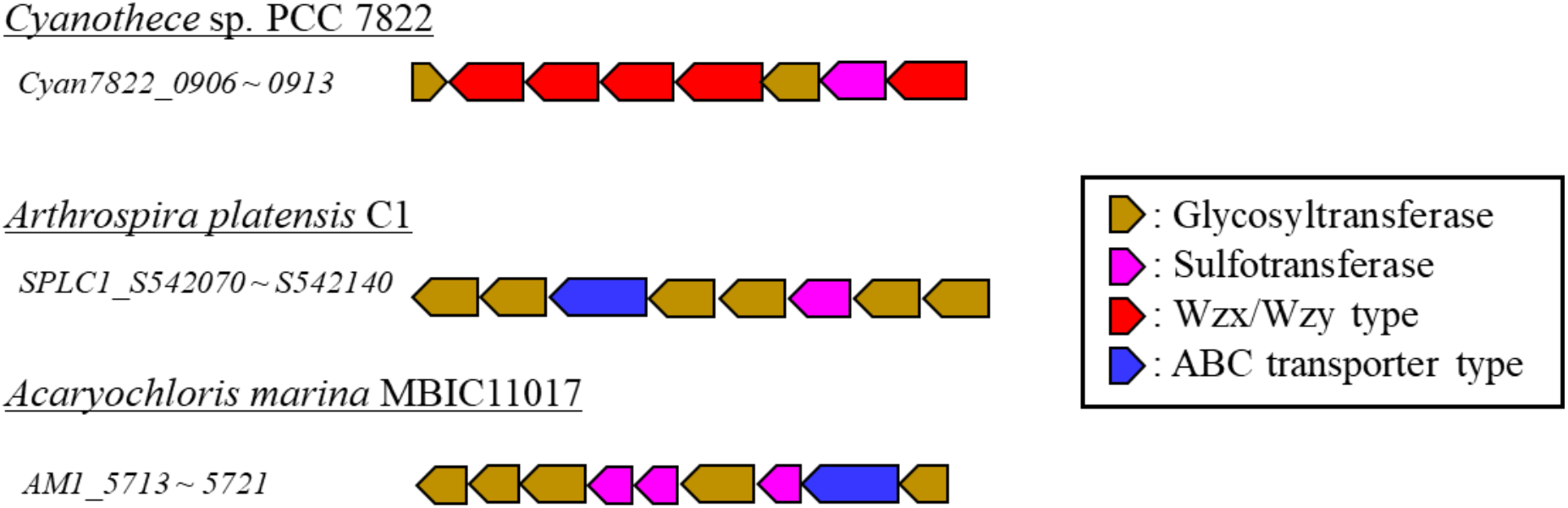
Typical examples of putative gene clusters for biosynthesis of sulfated polysaccharides in cyanobacteria. A part of each cluster harboring genes for sulfotransferases, glycosyltransferases, and polysaccharide biosynthesis/export systems (Wzx/Wzy type and ABC transporter type) in cyanobacterial genomes.

**Figure S9.**
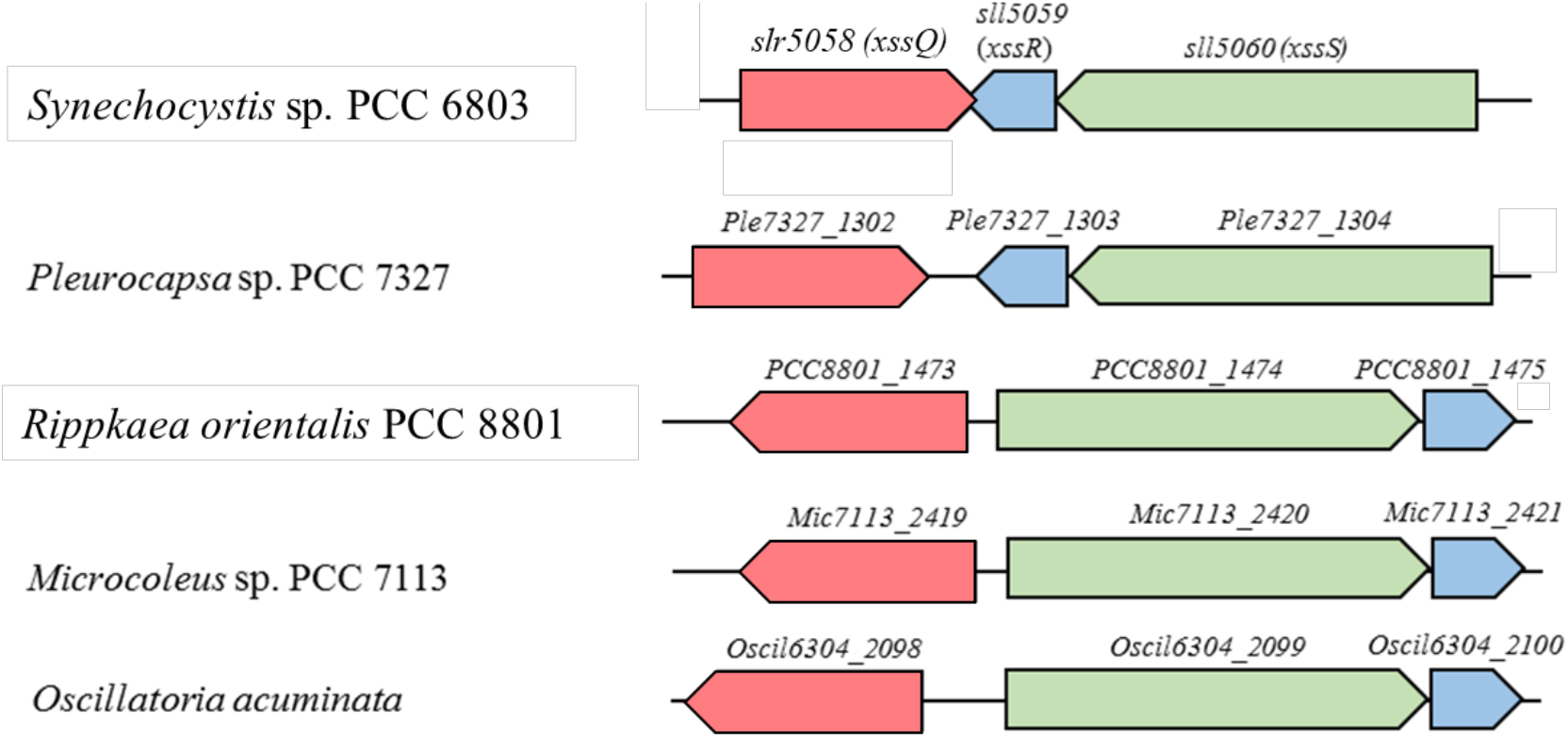
Homologous gene clusters of *xssS*/*xssR*/*xssQ* in some cyanobacteria. The colored blocks indicate genes with direction. The homologs of each gene are represented by their corresponding colors (red, *xssQ*; blue, *xssR*; green, *xssS*).

**Table S1.**
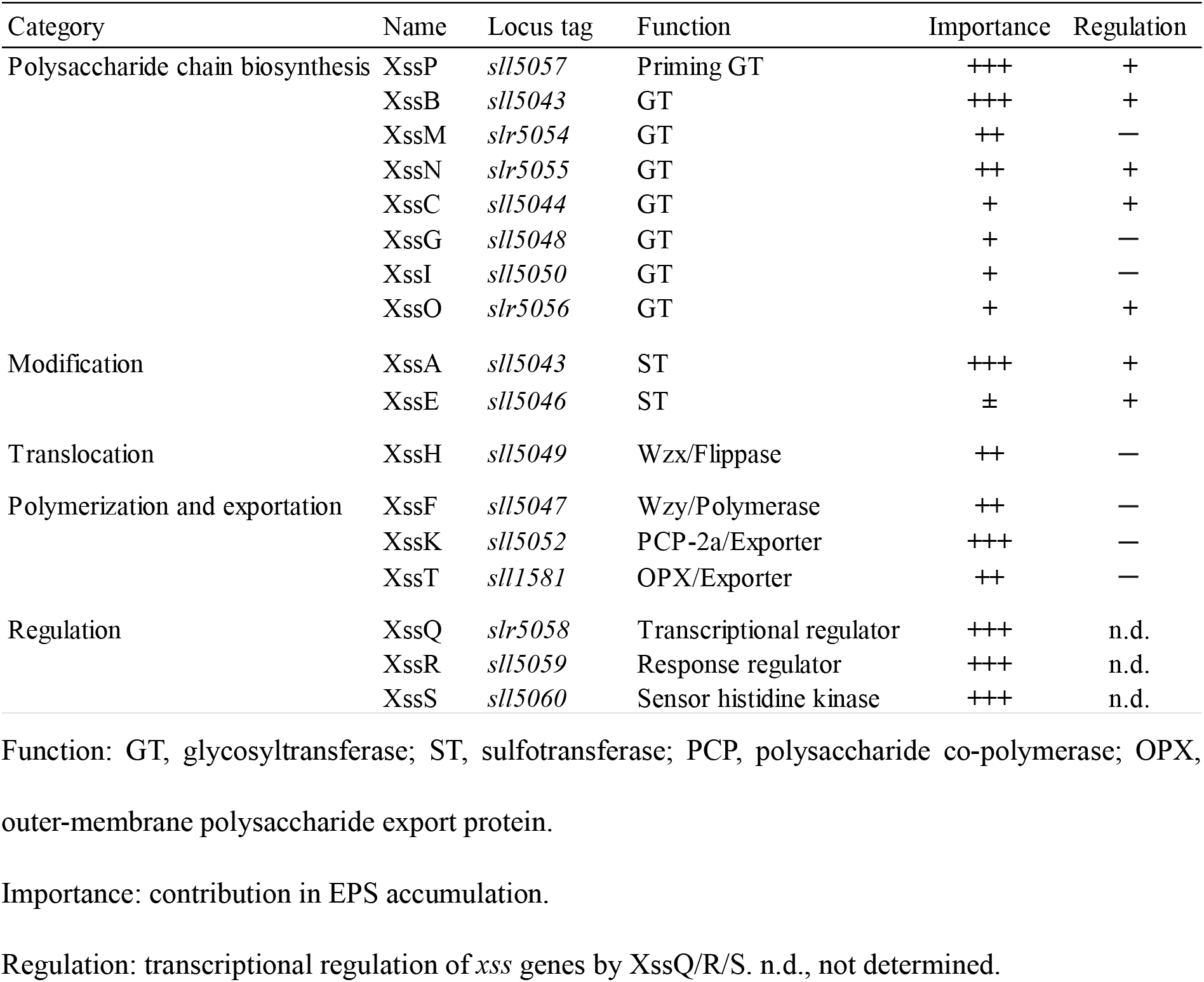
Summary of *Synechocystis* 6803 Xss proteins.

**Table S2.** EPS accumulation and bloom formation by WT *Synechocystis* 6803 and *xss* mutants.

| Strain name | Classification | Glc ($\mu\text{g}$) per $1.0 \times 10^8$ cells | Ratio (% WT) | Bloom |
| --- | --- | --- | --- | --- |
| WT(31°C) | | $0.526 \pm 0.062$ | 100.0 | + |
| WT(20°C) | | $1.629 \pm 0.472$ | 309.9 | + |
| $\Delta xssA$ | ST3 | $0.054 \pm 0.042$ | 10.4 | - |
| $\Delta xssB$ | GT2 | $0.040 \pm 0.029$ | 7.7 | - |
| $\Delta xssC$ | GT25 | $0.148 \pm 0.020$ | 28.1 | + |
| $\Delta xssE$ | ST3 | $0.402 \pm 0.127$ | 76.6* | + |
| $\Delta xssF$ | Wzy / polymerase | $0.107 \pm 0.008$ | 20.3 | - |
| $\Delta xssG$ | GT4 | $0.105 \pm 0.018$ | 20.1 | + |
| $\Delta xssH$ | Wzx / Flippase | $0.112 \pm 0.059$ | 23.0 | - |
| $\Delta xssI$ | GT4 | $0.113 \pm 0.044$ | 21.6 | + |
| $\Delta xssJ$ | - | $0.224 \pm 0.025$ | 42.6 | + |
| $\Delta xssK$ | PCP2a | $0.037 \pm 0.012$ | 7.0 | - |
| $\Delta xssM$ | GT2 | $0.049 \pm 0.025$ | 14.6 | - |
| $\Delta xssN$ | GT26 | $0.115 \pm 0.025$ | 21.8 | - |
| $\Delta xssO$ | GT2 | $0.161 \pm 0.025$ | 30.6 | + |
| $\Delta xssP$ | Priming GT | $0.039 \pm 0.010$ | 7.4 | - |
| $\Delta xssQ$ | Transcriptional regulator | $0.036 \pm 0.010$ | 6.8 | - |
| $\Delta xssR$ | Response regulator | $0.059 \pm 0.009$ | 11.2 | - |
| $\Delta xssS$ | Histidine kinase | $9.770 \pm 0.524$ | 1860 | ++ |
| $\Delta xssR-S$ | | $0.062 \pm 0.008$ | 11.8 | - |
| $\Delta xssK \sim M + \Delta xssS$ | | $0.044 \pm 0.013$ | 8.4 | - |
| OX-$xssR$ | | $10.426 \pm 0.225$ | 1984 | ++ |
| OX-$xssQ$ | | $7.359 \pm 2.36$ | 1400 | ++ |
| $\Delta xssQ + \text{OX-}xssR$ | | $0.077 \pm 0.027$ | 14.6 | - |
| $\Delta xssR + \text{OX-}xssQ$ | | $4.837 \pm 2.169$ | 920 | + |
| $\Delta sll1581$ ($xssT$) | OPX / exporter | $0.103 \pm 0.012$ | 19.6 | - |

Classification: ST, sulfotransferase; GT, glycosyltransferase; PCP, polysaccharide co-polymerase; OPX,
outer-membrane polysaccharide export protein. Total sugar content of the EPS fraction is expressed as
$\mu\text{g}$ glucose / $1 \times 10^8$ cells. The errors are based on SD ( $n = 3$ ). \*Close to the detection limit. EPS
accumulation ratio, percent of WT grown at 31°C. Bloom formation is summarized from Fig. 2.

**Table S3.**
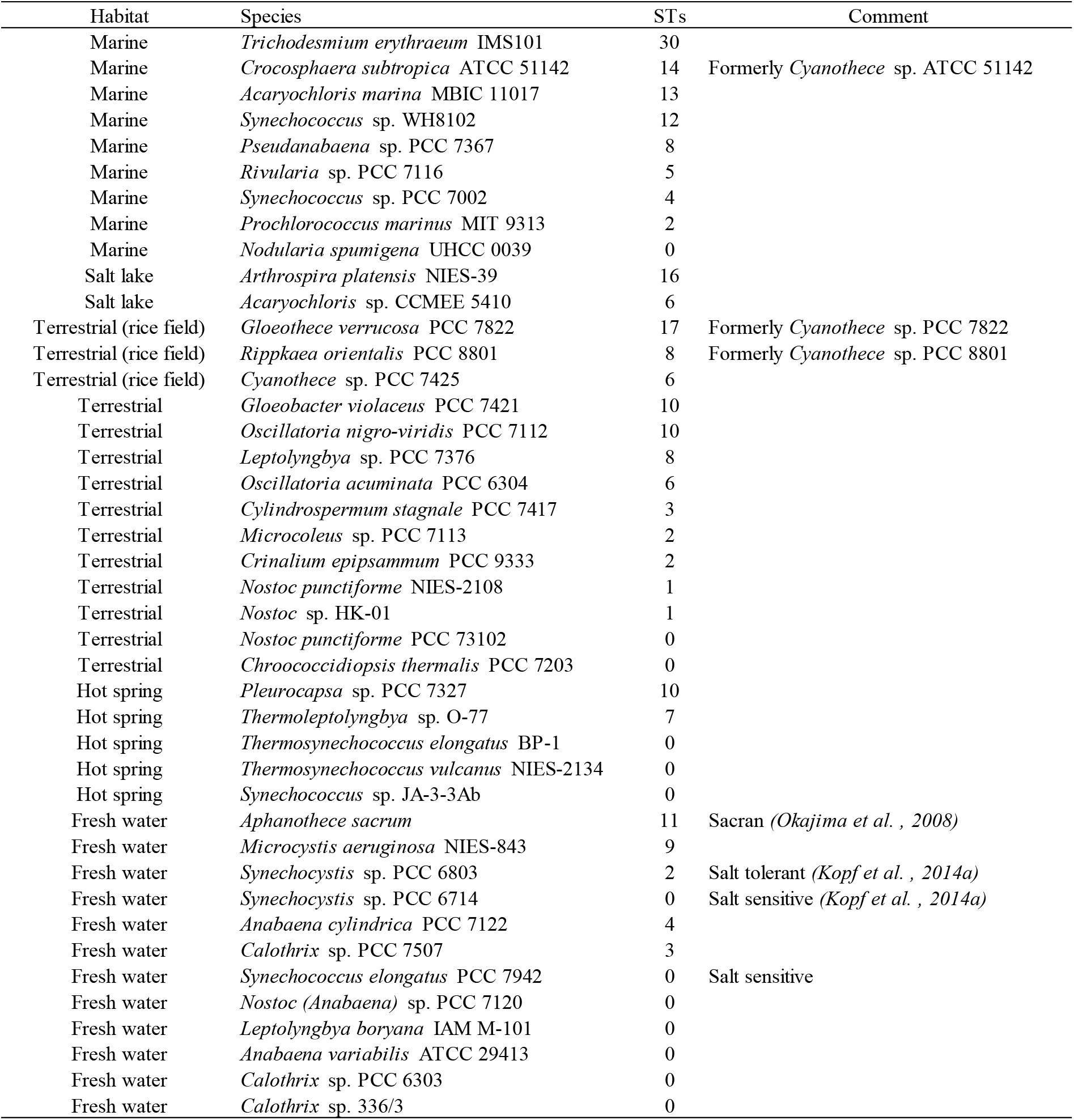
Number of sulfotransferase genes (STs) in the genome of cyanobacteria collected from various habitats.

**Table S4.**
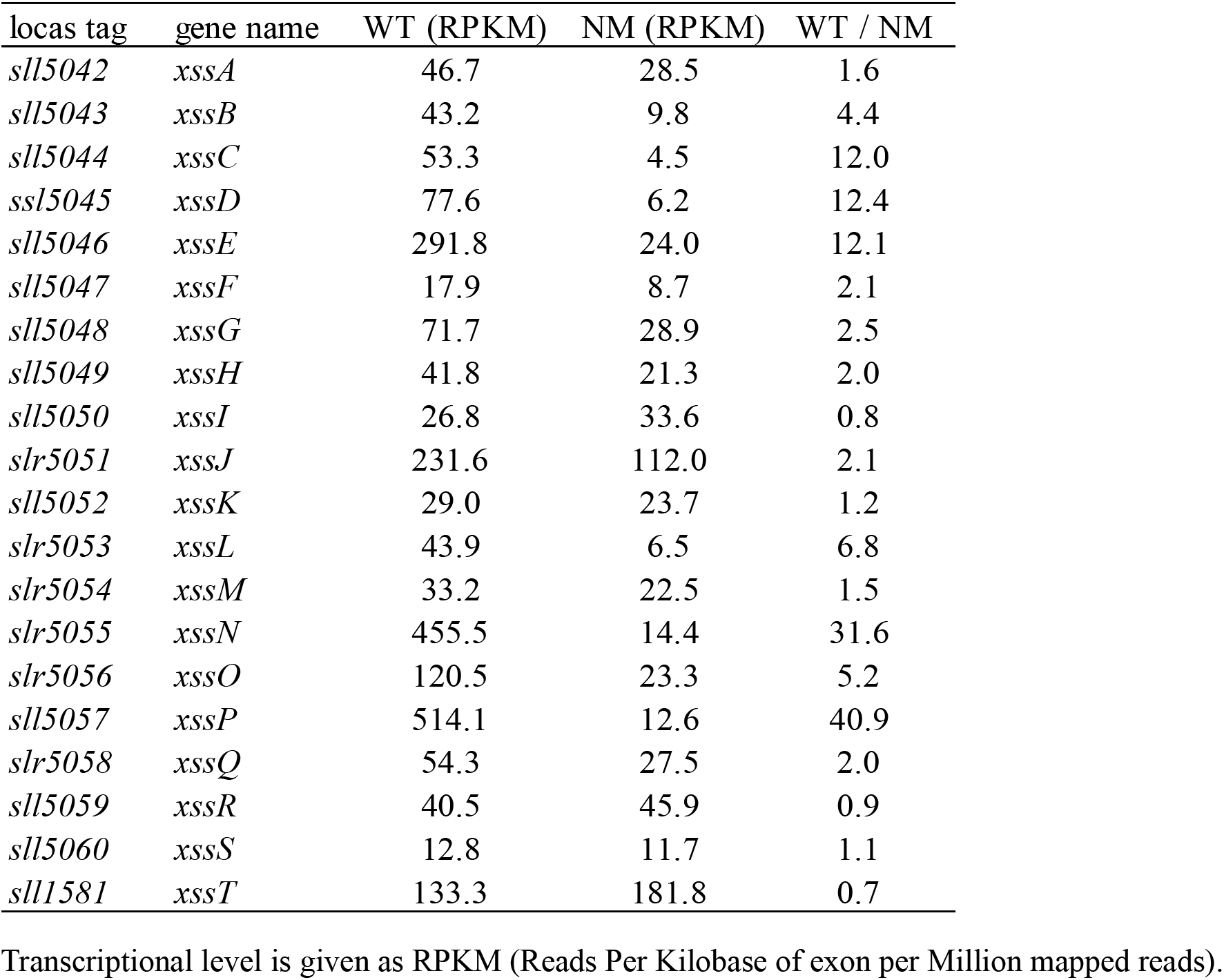
Transcriptional levels of *xss* genes in the motile (WT) and nonmotile (NM) substrains.

**Table S5.**
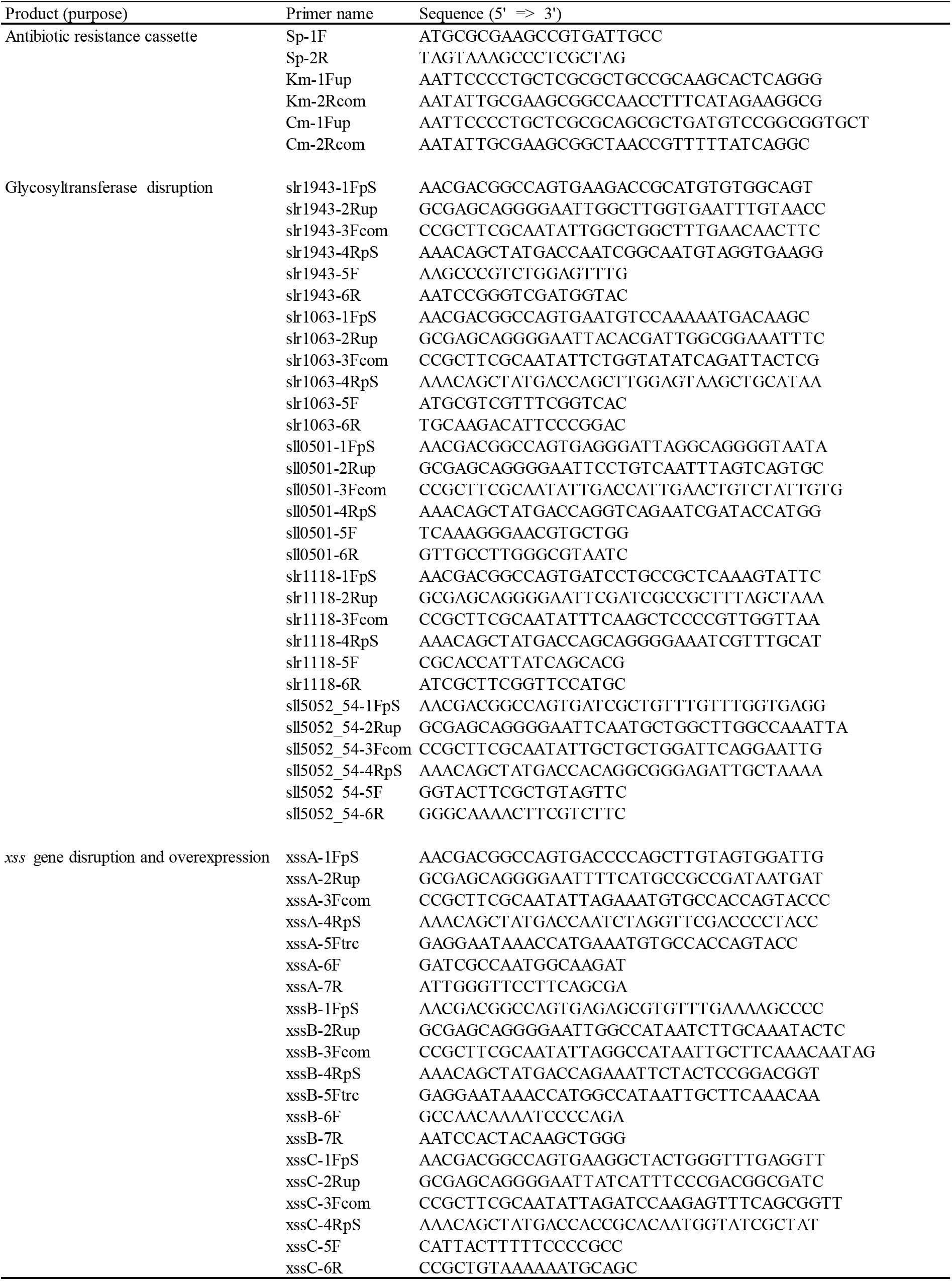
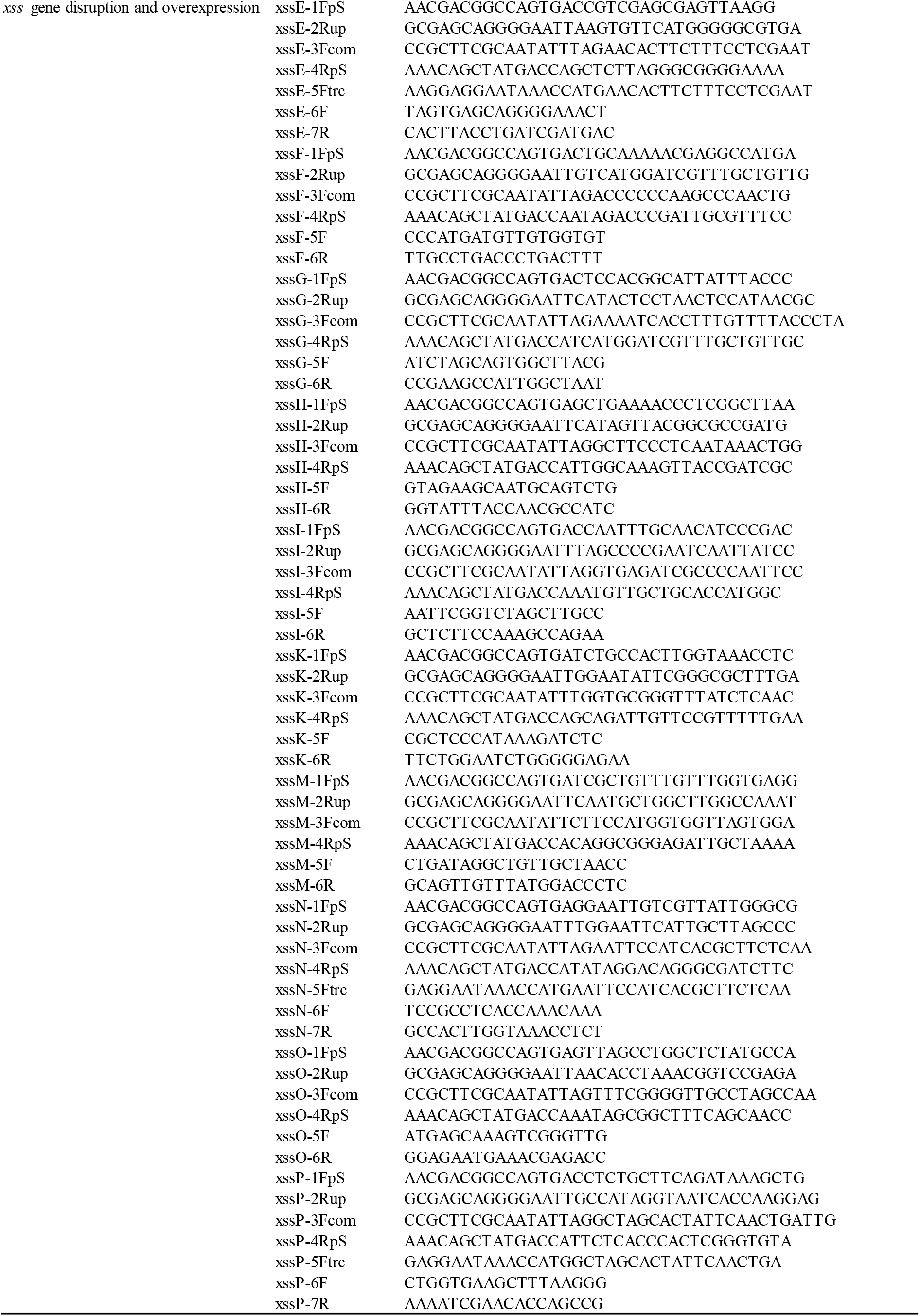
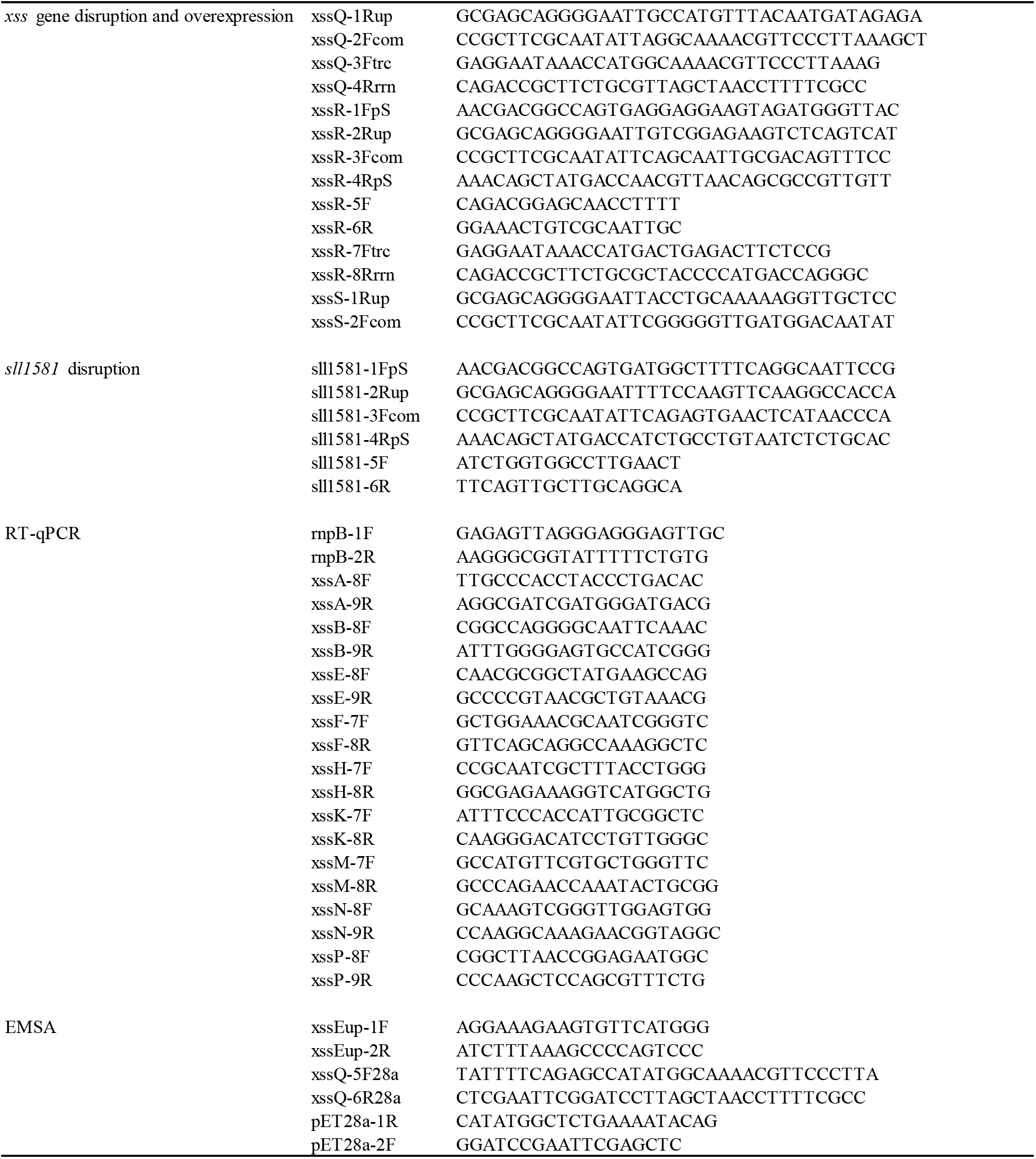
Primer sequences.

**Table S6.**
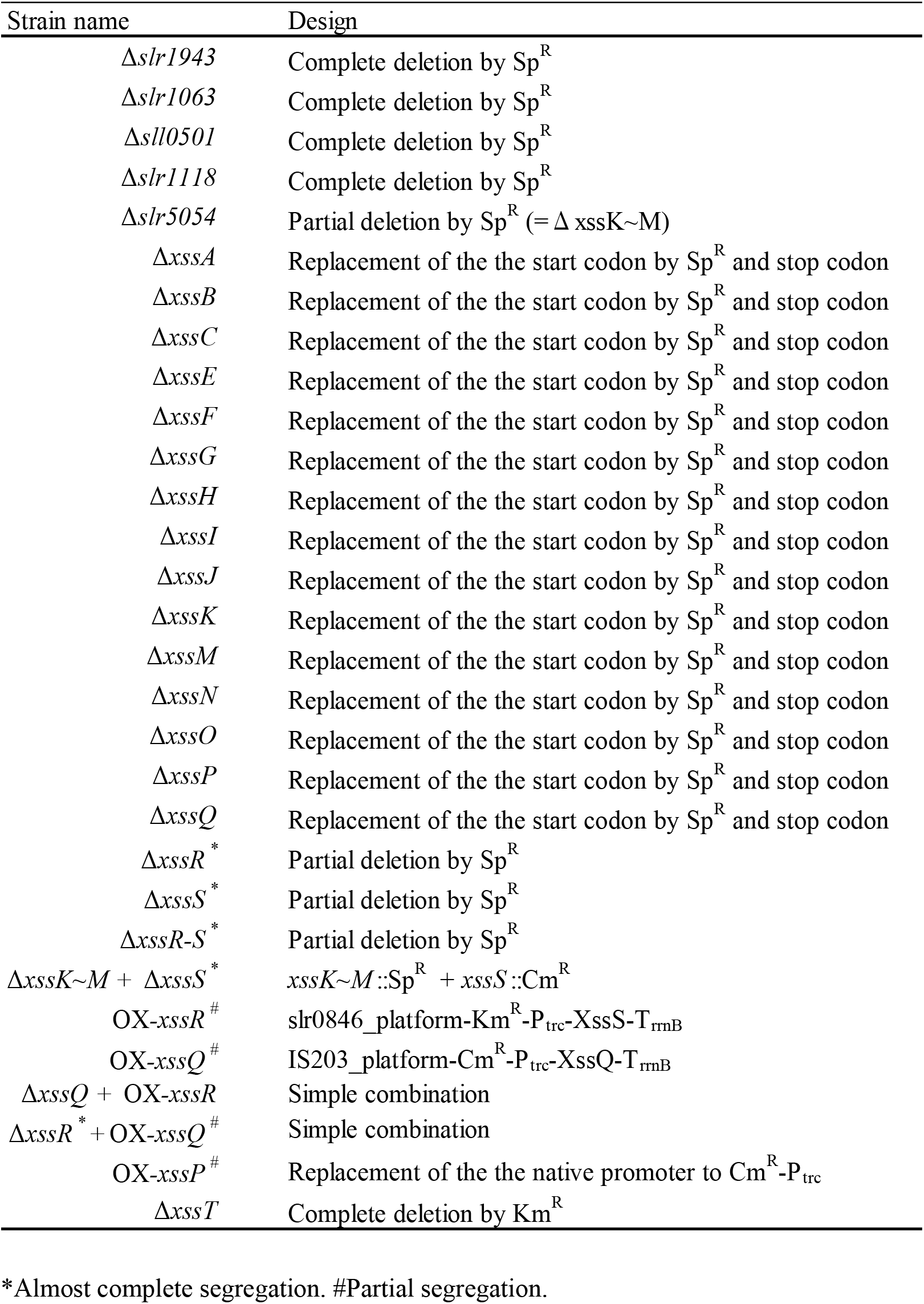
List of mutants.

